# Genetic studies of human-chimpanzee divergence using stem cell fusions

**DOI:** 10.1101/2021.09.24.461617

**Authors:** Janet H.T. Song, Rachel L. Grant, Veronica C. Behrens, Marek Kucka, Garrett A. Roberts Kingman, Volker Soltys, Yingguang Frank Chan, David M. Kingsley

## Abstract

Complete genome sequencing has identified millions of DNA changes that differ between humans and chimpanzees. Although a subset of these changes likely underlies important phenotypic differences between humans and chimpanzees, it is currently difficult to distinguish causal from incidental changes and to map specific phenotypes to particular genome locations. To facilitate further genetic study of human-chimpanzee divergence, we have generated human and chimpanzee auto-tetraploids and allo-tetraploids by fusing induced pluripotent stem cells (iPSCs) of each species. The resulting tetraploid iPSCs can be stably maintained and retain the ability to differentiate along ectoderm, mesoderm, and endoderm lineages. RNA sequencing identifies thousands of genes whose expression differs between humans and chimpanzees when assessed in single-species diploid or auto-tetraploid iPSCs. Analysis of gene expression patterns in inter-specific allo-tetraploid iPSCs shows that human-chimpanzee expression differences arise from substantial contributions of both *cis*-acting changes linked to the genes themselves, and *trans*-acting changes elsewhere in the genome. To enable further genetic mapping of species differences, we tested chemical treatments for stimulating genome-wide mitotic recombination between human and chimpanzee chromosomes, and CRISPR methods for inducing species-specific changes on particular chromosomes in allo-tetraploid cells. We successfully generated derivative cells with nested deletions or inter-specific recombination on the X chromosome. These studies identify a long distance *cis*-regulatory domain of the Fragile X-associated gene (*FMR1*), confirm an important role for the X chromosome in trans-regulation of other expression differences, and illustrate the potential of this system for more detailed mapping of the molecular basis of human and chimpanzee evolution.

**Significance Statement:** Comparative studies of humans and chimpanzees have revealed many anatomical, physiological, behavioral, and molecular differences. However, it has been challenging to map these differences to particular chromosome regions. Here, we develop a genetic approach in fused stem cell lines that makes it possible to map human-chimpanzee molecular and cellular differences to specific regions of the genome. We illustrate this approach by mapping chromosome regions responsible for species-specific gene expression differences in fused tetraploid cells. This approach is general, and could be used in the future to map the genomic changes that control many other human-chimpanzee differences in various cell types or organoids *in vitro*.

---

Humans have had a long-standing interest in the features that distinguish our species from other animals (1, 2). Comparative studies have characterized many morphological, physiological, and behavioral similarities and differences among great apes (3). Paleontological studies have traced the origin and timing of the appearance of various human features in the fossil record (4). More recently, advances in sequencing technologies have allowed for the comparative genomic analysis of humans, chimpanzees, other non-human primates, and even extinct archaic human lineages such as Neanderthals and Denisovans (5).

Whole genome comparisons indicate that approximately 4% of the base pairs in the human genome differ from those in chimpanzees. Sifting through this set of ~125 million DNA changes to separate the causal mutations contributing to phenotypic differences between humans and chimpanzees from inconsequential or neutral changes is a daunting problem, and has been compared to searching for needles in a haystack (3).

In evolutionary studies of other organisms, genetic crosses between different lineages have helped localize and prioritize chromosome regions that influence different traits. The formation of F1 hybrids, followed by chromosome recombination during meiosis, can be used to produce F2 offspring that inherit different combinations of alleles from the parental lineages. By comparing different genotypes and phenotypes across a large panel of meiotic mapping progeny, it has now been possible to map some evolutionary traits to particular chromosome regions in yeast, fruit flies, butterflies, sticklebacks, mice, and other organisms (6).

Traditional meiotic mapping approaches are limited to organisms that can be crossed to produce viable and fertile offspring. However, related approaches have also been developed for comparing genotypes and phenotypes in somatic cells without meiosis, when traditional crosses are not possible. Cells of even distantly related organisms can be fused *in vitro* to produce somatic cell hybrids that contain the genetic information from both lineages. The fused cells sometimes lose chromosomes of one or the other starting species, producing progeny cell lines that can be used to assign genes or cellular phenotypes to particular chromosomes (7). Hybrids can also be irradiated to fragment chromosomes and stimulate additional segregation of genetic information, an approach that has been used for fine mapping of genomic linkage relationships (8). Mitotic recombination within cultured cells can also be stimulated by mutations in DNA pathways, by chemicals that damage DNA, or by targeted breaks induced by Cas9 and guide RNAs designed to alter particular locations in the genome. Mutations and chemical inhibitors of the Bloom Syndrome helicase gene (*BLM*) have been used to recover homozygous mutants in somatic cell gene screens (9, 10) or to induce recombination between chromosomes of distantly related mouse strains for studies of the genomic basis of evolutionary differences (11). The ability to induce breaks at particular loci with CRISPR has also made it possible to choose both the location and the direction of recombination between genomes in non-meiotic cells, enabling high-resolution mapping without traditional crosses in yeast (12).

Development of similar approaches for human and chimpanzee cells would be very useful for studying the genomic basis of evolutionary differences that have evolved in hominids. Many molecular and cellular phenotypes that can be assayed and scored under cell culture conditions are known to differ between humans and chimpanzees. Recent studies have generated well-matched sets of human and chimpanzee induced pluripotent stem cell (iPSC) lines (13), and have shown that human and chimpanzee iPSCs can be fused to produce hybrids useful for comparing species-specific expression in cortical spheroids and neural crest cells (14, 15). Here we generate both auto-tetraploid (same species) and allo-tetraploid (different species) fusion lines from human and chimpanzee iPSCs, and use them to identify whether gene expression differences are due to *cis*- or *trans*-acting differences between species. We also test both random and targeted methods for stimulating DNA breaks and chromosome exchanges in allo-tetraploid iPSCs, providing a general method for further localizing the specific genomic changes that underlie human and chimpanzee differences *in vitro*.

## Results

### Generation and initial characterization of auto- and allo-te-traploid iPSC lines

To generate auto- and allo-tetraploids, we labeled human and chimpanzee iPSC lines (13) with diffusible fluorescent dyes and fused them using electrofusion (Fig. 1A, Materials and Methods). Tetraploid cells were enriched by either fluorescence-activated cell sorting (FACS) or manual inspection and grown clonally. Successful fusion in expanded clones was confirmed by FACS analysis for DNA content using propidium iodide and by karyotyping. In total, we generated 2 human auto-tetraploid lines (“H1H1” lines, from human iPSC line H23555); 5 chimpanzee auto-tetraploid lines (“C1C1” lines from chimpanzee iPSC line C3649); and 22 human-chimpanzee allo-tetraploid lines from different fusion events including 12 “H1C1” lines derived from H1 and C1 and 10 “H2C2” lines derived from human iPSC line H20961 (H2) and chimpanzee iPSC line C8861 (C2) (Table S1).

**Fig. 1.**
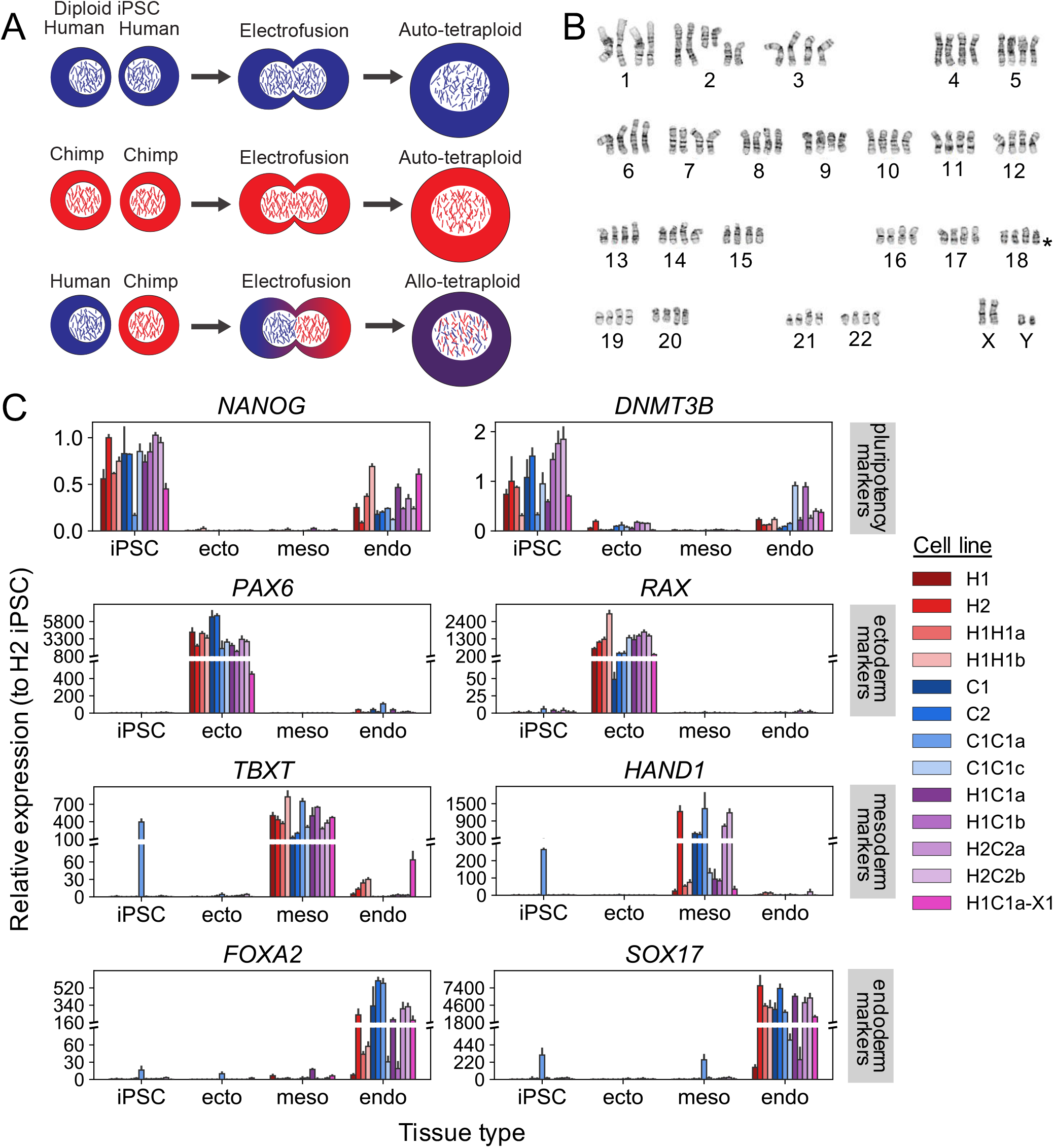
Generation and differentiation of auto- and allo-tetraploid iPSC lines. Auto- and allo-tetraploid cells contain the expected number of chromosomes and express expected marker genes after trilineage differentiation. **(A)** Human and chimpanzee diploid iPSC lines were labeled with diffusible dyes and subjected to electrofusion to generate auto- and allo-tetraploid iPSC lines. **(B)** Tetraploid lines (H2C2a shown) exhibit karyotypes with 4 copies of each chromosome. Asterisk denotes location of the common iPSC human chr18q deletion (16), present in a subset of our cell lines. See Table S1 for detailed karyotype description of all lines. **(C)** Relative expression of pluripotency (*NANOG, DNMT3B*), ectoderm (*PAX6, RAX*), mesoderm (*TBXT, HAND1*), and endoderm (*FOXA2, SOX17*) marker genes tested via RT-qPCR after incubating cell lines under trilineage differentiation conditions. Cell lines tested: two human diploid lines (H1, H2), two human auto-tetraploid lines (H1H1a, H1H1b), two chimpanzee diploid lines (C1, C2), two chimpanzee auto-tetraploid lines (C1C1a, C1C1c), four allo-tetraploid lines (H1C1a, H1C1b, H2C2a, H2C2b), and one fluorescently-marked allo-tetraploid line (H1C1a-X1). Gene expression is plotted relative to a human diploid undifferentiated iPSC line (H2). Error bars represent the standard deviation of *N* = 3 cell culture replicates maintained as iPSCs or differentiated independently. 146 of 156 gene expression differences between undifferentiated cells and the tissue type in which a marker is expected to be expressed are significant by two-tailed Student’s t-test at 5% FDR (see Table S3 for complete *p*-value list).

Tetraploid iPSCs were larger than diploid cells but had normal morphology and could be routinely propagated under the same conditions as diploid iPSCs (Fig. S1). We performed G-banded karyotyping on the initial diploid parental lines, as well as the newly generated auto- and allo-tetraploid lines to examine their genome stability (Table S1). Fusion lines showed the tetraploid karyotypes expected from fusing their originating diploid lines. However, some of the tetraploid lines contained additional chromosomal abnormalities, including aneuploidies common to diploid human iPSC cultures (16) such as deletion of human chr18q (asterisk in Fig. 1B).

To assess the pluripotency and differentiation potential of the tetraploid iPSC lines, we differentiated representative diploid (H1, H2, C1, C2), auto-tetraploid (H1H1a, H1H1b, C1C1a, C1C1c), and allo-tetraploid (H1C1a, H1C1b, H2C2a, H2C2b) lines into ectoderm, mesoderm, and endoderm (Materials and Methods). Quantitative PCR for the expression of pluripotency (*NANOG, DNMT3B*), ectoderm (*PAX6, RAX*), mesoderm (*TBXT*, *HAND1*), and endoderm (*FOXA2, SOX17*) markers showed specific differentiation of tetraploid lines into all three lineages (Fig. 1C, Table S3, SI Methods). For endoderm differentiation, a subset of lines (H1, C1C1c, H1C1b) showed lower expression of endoderm markers compared to all other cell lines as well as persistent expression of pluripotency marker genes. Tetraploid cells thus retain broad differentiation abilities, but conditions may need to be optimized for particular cell lines or differentiation endpoints.

### Diploid and auto-tetraploid iPSC lines have similar gene expression profiles

To examine whether tetraploidization altered normal gene expression patterns, we used RNA sequencing (RNAseq) to characterize transcriptional differences due to ploidy, but not to species differences (i.e. H1 vs. H1H1 and C1 vs. C1C1). At a false discovery rate (FDR) of 5%, we detected 189 differentially expressed genes between H1 and H1H1, and 181 differentially expressed genes between C1 and C1C1 with at least a 2-fold change in expression (Table S4, SI Methods). Neither set of differentially expressed genes was enriched for gene ontology categories (SI Methods), and only 13 genes were differentially expressed in both H1 compared to H1H1 and C1 compared to C1C1. We conclude that the creation of tetraploid cells alone does not activate a coordinated set of gene expression changes.

To assess gene expression variability between different cell lines from the same species, we also profiled global RNA patterns from a second set of human and chimpanzee diploid iPSC lines. We detected 410 differentially expressed genes between H1 and H2 and 181 differentially expressed genes between C1 and C2 at an FDR of 5% with at least a 2-fold change in expression (Table S4). Using principal component analysis, we found that global transcriptional profiles grouped by species, with human-derived lines clustering separately from chimpanzee-derived lines, and that diploid lines clustered more closely with their derived auto-tetraploid line than another diploid line of the same species (Fig. 2A). These results indicated that the transcriptional profiles of the diploid lines and their derived auto-tetraploid lines were at least as similar as the transcriptional profiles of two diploid lines from the same species. Taken together, our data suggest that tetraploid iPSCs behave similarly to diploid iPSCs at the level of gene expression.

**Fig. 2.**
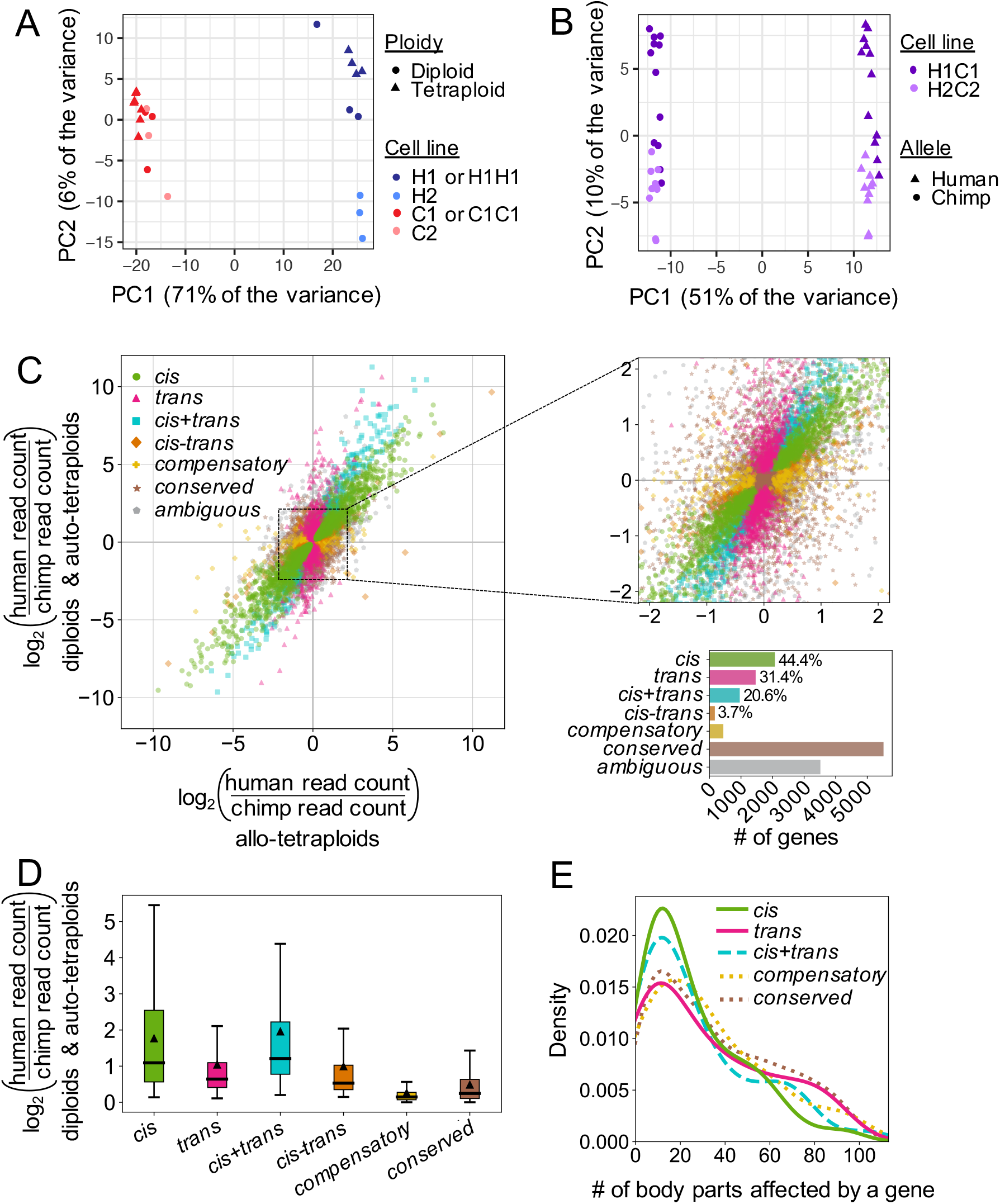
Gene expression profiling of human and chimpanzee diploid, auto- and allo-tetraploid iPSC lines. Tetraploidization does not result in coordinated gene expression changes, but thousands of genes are expressed differently between human and chimpanzee iPSCs due to a mixture of *cis*- and *trans*-regulatory changes. **(A)** Principal component analysis (PCA) of RNA sequencing of H1, H2, C1, C2, H1H1, and C1C1 diploid and auto-tetraploid iPSC lines. The cell lines cluster by species along PC1 and by cell line along PC2. Auto-tetraploid lines cluster with their cognate diploid line. **(B)** PCA of RNA sequencing of H1C1 and H2C2 allo-tetraploid lines. Allo-tetraploid lines are each represented by two dots, one for reads mapping to the human transcriptome and one for reads mapping to the chimpanzee transcriptome (Materials and Methods). Expression from human alleles (triangles) cluster separately from chimpanzee alleles (circles) in allo-tetraploid lines along PC1. PC2 separates the two sets of allo-tetraploid cell lines. **(C)** Each gene’s expression pattern was classified by regulatory type (*cis*, *trans, cis+trans, cis-trans, compensatory, conserved*, or *ambiguous*) by comparing differential gene expression between human- and chimpanzee-only iPSCs (x-axis) and allele-specific gene expression between human and chimpanzee alleles within allo-tetraploid iPSCs (y-axis). Left: data for all genes. Upper right: zoom-in of dense center region. Lower right: bar graph indicating number of genes per category and relative contribution (percentage) of each category to genes with human-chimpanzee regulatory differences. **(D)** Box plot showing distribution of effect sizes for gene expression changes in each regulatory category. Median effect size is indicated by thick horizontal lines, and mean effect size is indicated by triangles. All pairwise comparisons are statistically significant (adjusted *p* < 0.012 by two-tailed Mann-Whitney U test). **(E)** Density plot (smoothed histogram) showing the distribution of body parts influenced by genes (according to the Gene ORGANizer database (19)) in each regulatory category. For genes classified as *cis*, *trans*, and *cis*+*trans*, only genes with |*log*_2_(*FC*)|≥1 are plotted. The *cis-trans* category is not included because only 5 genes have |*log*_2_(*FC*)|≥1. Note that genes classified as *cis* or *cis+trans* tend to influence fewer body parts than *conserved* genes (median 18 body parts for both *cis* and *cis+trans* genes compared to median 30 body parts for *conserved* genes, adjusted *p* = 0.00028 and *p* = 0.0035 by two-tailed Mann-Whitney U test after FDR correction). This trend is not observed for *trans* and *compensatory* regulatory types (median 24 and 27 body parts, adjusted *p* = 0.11 and *p* = 0.21, respectively).

### Differential gene expression and allele-specific gene expression reveal human- and chimpanzee-specific gene expression profiles

We next used our RNAseq data to identify gene expression differences between human and chimpanzee iPSCs (Table S5). Differential gene expression (DE) analysis between human-only and chimpanzee-only iPSC lines identified 5,984 genes differentially expressed between species. There were no significant gene ontology enrichments for DE genes with at least a 2-fold change in expression (SI Methods). Allelespecific expression (ASE) comparisons between the human allele and the chimpanzee allele in allo-tetraploid iPSC lines identified 4,540 allele-specifically expressed genes. ASE results from this study and the ASE results from a previous study (14) that independently generated human-chimpanzee allo-tetraploid fusions from similar diploid iPSC lines were highly concordant (Pearson’s *r* = 0.72; Fig. S2), suggesting that human-chimpanzee gene expression differences are robust and reproducible across laboratories.

### *Cis-* and trans-acting regulatory changes are both important contributors to human-chimpanzee gene expression differences

Determining whether gene expression differences between two species are due to *cis*-acting or *trans*-acting regulatory changes is possible when gene expression can be compared between each single-species and a hybrid (17). We therefore leveraged the RNAseq data from human-only, chimpanzee-only, and human-chimpanzee allo-tetraploid iPSC lines to determine the regulatory type of genes that were differentially expressed between human-only and chimpanzee-only iPSCs (Materials and Methods). Specifically, when a *cis*-acting regulatory change causes a gene to be differentially expressed, the expression difference should be maintained in allo-tetraploid cells where both human and chimpanzee alleles are in the same *trans*-acting environment. Conversely, when a *trans*-acting regulatory change causes a gene to be differentially expressed, the expression difference should disappear in allo-tetraploid cell lines.

Our regulatory type classifications identified 5,956 genes with no net regulatory changes between our human-only and chimpanzee-only iPSC lines. Of these, 92.6% (5,515 genes) were classified as *conserved* between human and chimpanzee, and 7.4% (441 genes) were classified as *compensatory* (*cis* and *trans*-regulatory differences acting in opposite directions resulting in no net expression difference between species) (Fig. 2C).

Of 4,671 genes with regulatory changes between human-only and chimpanzee-only iPSC lines, 44.4% (2,073 genes) were regulated primarily in *cis*, 31.4% (1,465 genes) were regulated primarily in *trans*, and the remaining 1,133 genes were regulated both in *cis* and in *trans* (Fig. 2C). This final category was further broken down into a *cis+trans* category (*cis*- and *trans*-regulatory changes acting in the same direction) and a *cis-trans* category (*cis-* and *trans*-regulatory changes acting in opposite directions). This yielded 20.6% (961 genes) and 3.7% (172 genes) regulated in *cis+trans* and *cis-trans*, respectively. Other genes that did not satisfy the conditions for any category (3,515 genes) were classified as *ambiguous*.

Genes with primarily *cis*-regulatory changes had a larger median effect size than genes with primarily *trans*-regulatory changes (median |*log*_2_(*FC*)| of 1.09 vs. 0.64, *p* < 10^−56^ by 2-tailed Mann-Whitney U test; Fig. 2D). Genes classified as *cis+trans* had the highest effect size of any regulatory type category (median |*log*_2_(*FC*)| of 1.21).

Gene ontology enrichments for genes classified as *trans* included processes related to the skeletal, cartilage, and muscular systems (Table S6), suggesting that some of the dramatic skeletal and muscular differences between humans and chimpanzees may be driven by *trans*-acting regulatory changes. Enrichments for genes classified as *conserved* were related to voltage-gated ion channels (Table S6), which are important for maintaining critical features of iPSCs including proliferation capacity and differentiation potential (18). Finally, genes classified as *compensatory* had enrichments related to ligase activity, neurexin protein binding, and phosphatidylserine binding, while all other regulatory type classifications had no significant gene ontology enrichments.

We also used the Gene ORGANizer database (19), which links genes to the body parts they affect based on phenotypes associated with Mendelian disorders, to test whether genes that were differentially expressed between humans and chimpanzees tend to influence more or fewer biological systems than *conserved* genes. We found that genes with primarily *cis*-regulatory changes and at least a 2-fold change in expression influenced a median of 18 body parts compared to a median of 30 body parts influenced by *conserved* genes (adjusted *p* = 2.8*x*10^−4^ by two-tailed Mann-Whitney U test; Fig. 2E). Interestingly, the greater the expression differences between human and chimpanzee as indicated by higher |*log*_2_(*FC*)|, the fewer body parts a gene with primarily *cis*-regulatory changes tended to influence (Fig. S3). Similar trends were observed for genes classified as *cis*+*trans* but not other regulatory categories.

Removing reads mapping to genes on chromosomes that were karyotypically abnormal in any of our iPSC lines did not significantly change our regulatory type classification or effect size results (Fig. S4). Together, our results indicate that both *cis-* and *trans*-acting regulatory changes are important contributors to the widespread gene expression differences between humans and chimpanzees in iPSCs, with *cis*-regulatory changes tending to be larger and to act on genes affecting fewer biological systems (Fig. 2C-E).

### Prospects for genetic mapping

Further localization of both *cis*- and *trans*-regulatory differences would be greatly aided if it were possible to generate mapping panels that carry different dosages of human and chimpanzee alleles at known locations throughout the genome. Previous studies in yeast, *Drosophila*, and cultured mammalian cell lines have used mitotic recombination to generate useful mapping panels from somatic cells (11, 12, 20, 21). To boost the rate of mitotic recombination, common strategies have been to treat cells with small molecules that promote DNA damage (22, 23), or to induce targeted recombination at specific loci using CRISPR/Cas9 (12, 20).

To assess whether small molecules could stimulate mitotic crossovers in iPSCs, we performed sister chromatid exchange (SCE) assays by incubating cells with BrdU for two cell cycles (24). Chromosomes where both strands incorporate BrdU stain lighter than chromosomes where only one strand has incorporated BrdU, making it possible to visualize SCE events in mitotic chromosome spreads (Fig. 3A). We tested camptothecin, a topoisomerase inhibitor previously found to induce SCE events in iPSCs (22). Consistent with prior findings, treatment of 100nM camptothecin for 1 hour induced a 4.5-fold increase in SCE events in both auto- and allo-tetraploid iPSCs (*p* < 10^−8^ by one-tailed Student’s t-test; Fig. 3B).

**Fig. 3.**
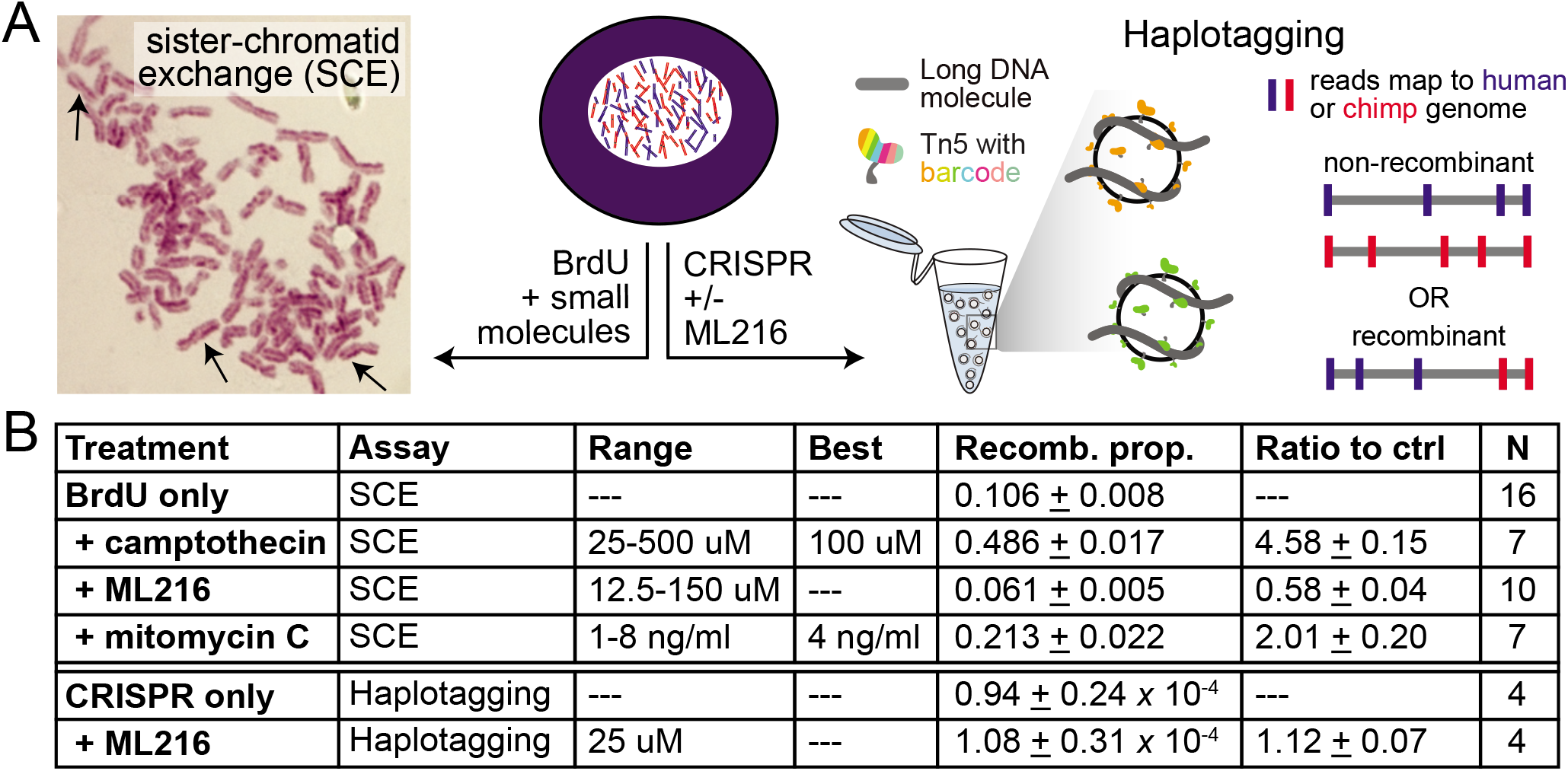
Effect of small molecules on chromosome recombination frequencies in allo-tetraploid cells. The small molecules camptothecin, ML216, and mitomycin C were assessed for their effect on intra- and inter-specific recombination. **(A)** Allo-tetraploid cells were treated with BrdU and small molecules to measure intra-specific sister chromatid exchange events by microscopy (left), or treated with Cas9 and guide RNAs with or without ML216 followed by haplotagging to identify inter-specific recombinant molecules by sequencing (right) (Materials and Methods). **(B)** The proportion of chromosomes that had SCE events after treatment with the indicated concentrations of each small molecule is listed (recomb. prop.). N: number of replicate experiments where the recomb. prop. was quantified for all concentrations or the best concentration (when indicated). Note BrdU was added to all cells to visualize SCE, and all BrdU+drug treatments were compared to the BrdU-only condition (ratio to ctrl). For haplotagging, the effect of CRISPR guides targeting specific loci was assessed with or without ML216. Compared to CRISPR alone, ML216 may elevate the proportion of inter-specific recombinant molecules genome-wide (ratio to ctrl). Values are mean + SEM.

We also tested ML216, an inhibitor of the Bloom syndrome helicase (BLM), which has been found to induce SCE events in cultured human cells (9, 10, 23). However, we found that treatment with ML216 over a range of concentrations from 12.5*μ*M to 150*μ*M did not increase the rate of SCE events in iPSCs. We additionally tested mitomycin C, which crosslinks DNA and is known to induce SCE events in yeast and fungi (25). Treatment of 4ng/ml mitomycin C for 24 hours in tetraploid iPSCs increased the rate of SCE events by 2-fold (*p* < 10^−5^ by one-tailed Student’s t-test; Fig. 3B). Although SCE assays can only reliably assess intra-specific crossover events, these results suggest that the application of camptothecin or mitomycin C to allo-tetraploid iPSCs has the potential to similarly increase the rate of inter-specific mitotic recombination.

An alternate approach is to induce targeted crossovers using CRISPR/Cas9. This strategy has previously been used to induce recombination in yeast and *Drosophila* (12, 20). To determine the rate of inter-specific recombination events at target loci, we used a recently developed technique called haplotagging to directly detect recombinant junctions by barcoding DNA molecules prior to sequencing (26, 27). Following sequencing, reads were aligned to a composite human-chimpanzee genome and comparatively assigned to their species of origin. Reads derived from the same DNA molecule were tagged with the same barcode, enabling molecule reconstruction (Materials and Methods). Barcoded molecules that mapped to orthologous intervals in human and chimpanzee and showed switched runs of variants from one species to the other (human-to-chimp or vice versa), were scored as likely inter-specific recombination events within the corresponding genomic interval.

In the allo-tetraploid line H1C1a, we targeted genomic loci on chr20q13.33, chr21q22.3, and chrXq28 with CRISPR guide RNAs (gRNAs) and then performed haplotagging to over 200x molecular coverage (Fig. S5, Fig. S6). Based on a recent study suggesting that ML216 acts synergistically with CRISPR/Cas9 to induce loss of heterozygosity at targeted loci in human iPSCs (21), we also assessed whether the addition of 25*μ*M of the BLM inhibitor ML216 starting 12 hours before gRNA targeting and ending 48 hours posttargeting would affect the rate of inter-specific recombination. We did not observe an enrichment in inter-specific recombination events at any of the target loci with or without ML216 treatment (Fig. S6). However, genome-wide inter-specific recombination events trended 1.12-fold higher when comparing ML216-treated samples against samples that were only treated with CRISPR/Cas9 (*p* = 0.088 by one-tailed Student’s t-test; Fig. 3B). With ML216 treatment, we detected a total of 43,256 inter-specific recombination events in ~389 million analyzed molecules. Further investigation will be required to assess whether ML216 significantly increases the rate of inter-specific recombination and whether other small molecules such as camptothecin can also increase the rate of recombination events between human and chimpanzee chromosomes in allo-tetraploid iPSCs.

### Targeted *cis*- and *trans*-mapping on the X chromosome

To further enrich for cells that may contain inter-specific mitotic recombination events at specific loci, we fluorescently marked allo-tetraploid cells at distal chromosome ends. This allowed us to use fluorescence-activated cell sorting (FACS) to isolate cells with expected signatures of recombination (Fig. 4A). The gene density and enrichment of disease-related genes on distal chrX, particularly chrXq28, made the distal region of chrX a particularly attractive target for further study (28). Through two rounds of CRISPR/Cas9-mediated homologous recombination (HR), we generated five allo-tetraploid lines derived from H1C1a and one from H2C2b, each carrying GFP on the human chrX and mCherry on the chimpanzee chrX (Fig. S7). Some allo-tetraploid cells that underwent two rounds of CRISPR/Cas9 HR-insertion maintained largely normal karyotypes while others showed more extensive aneuploidies (Table S1).

**Fig. 4.**
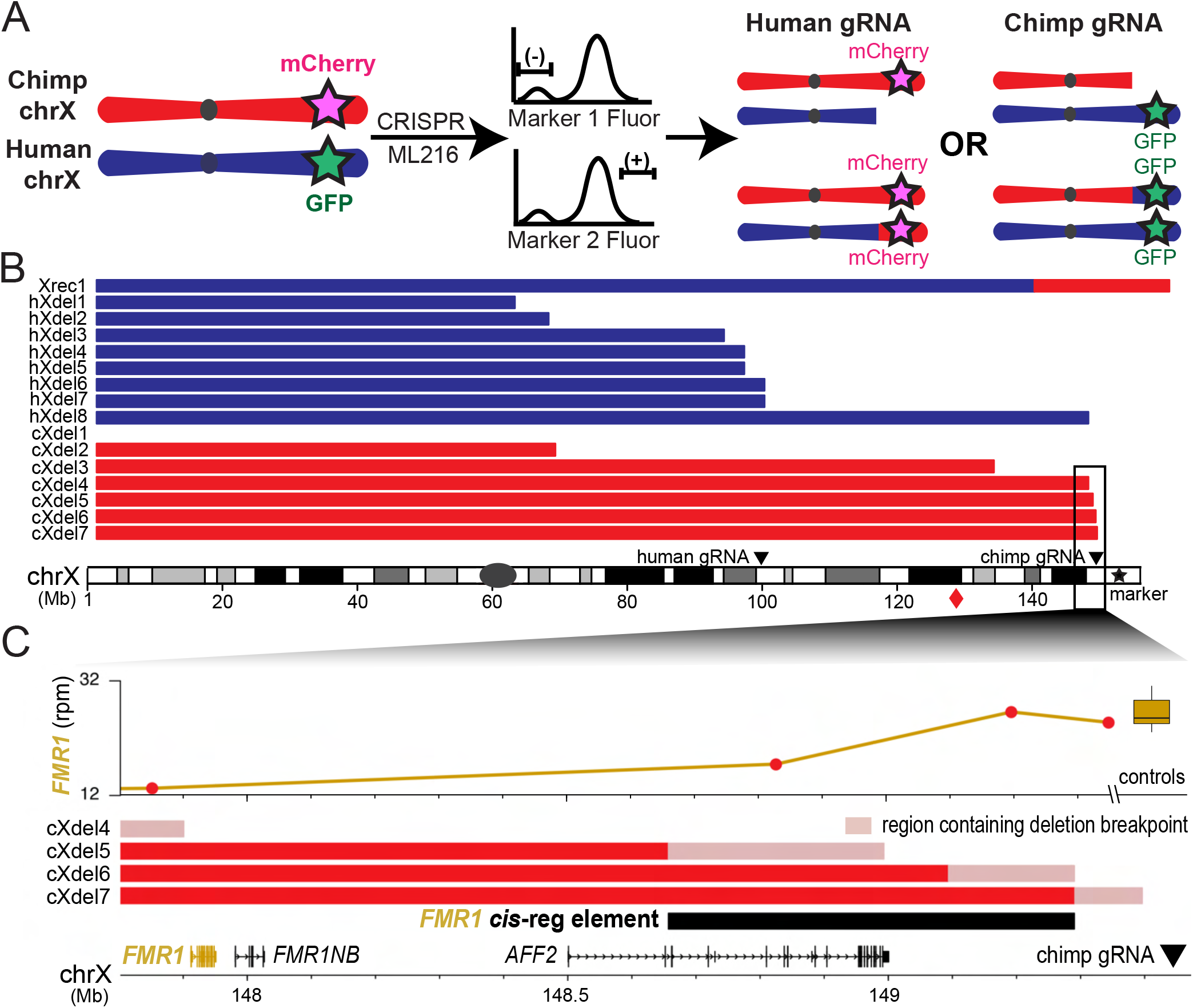
X chromosome targeting generates recombinant and deletion lines for genetic mapping. Cell lines with recombination and deletion events on the X chromosome were isolated by FACS and used for further mapping of regulatory sequences. **(A)** Allo-tetraploid cells marked with GFP on the distal human chrX and mCherry on the distal chimpanzee chrX were treated with ML216, Cas9, and species-specific chrX guide RNAs, followed by FACS for expected signatures of deletion or recombination (loss of fluorescent marker on targeted chrX, retention or gain of fluorescent marker from homologous, untargeted chrX). **(B)** Cell lines recovered from sorting contained either a recombinant chrX or species-specific distal chrX deletions. Breakpoint locations for the recombinant (Xrec1), human deletions (hXdel#), and chimpanzee deletions (cXdel#) are shown relative to human chrX. Positions of species-specific guide RNAs are shown. Diamond symbol indicates the distal portion of chimpanzee chrX that is recombined onto the first 140.1Mb of human chrX in Xrec1. **(C)** Level of *FMR1* expression drops in cells with loss of a distal downstream region from the chimpanzee chrX. Although line cXdel5 does not delete the chimpanzee allele of *FMR1*, it shows reduced *FMR1* expression compared to two further downstream deletions (cXdel6 and cXdel7) and the control lines (boxplot). Red: regions of chrX still present in lines. Pink: regions containing deletion breakpoints based on PCR assays and gene expression profiling (Materials and Methods).

Because the allo-tetraploids were derived from fusions of male cells, only one X chromosome was present from each species. In the double fluorescently-marked lines, cells with no recombination events on chrX should carry both human and chimpanzee fluorescent markers, while recombinant cells should carry two copies of a single marker from either human or chimpanzee chrX. We treated the double fluorescently-marked allo-tetraploid line H1C1a-X1 with either a chimpanzee-specific gRNA targeting chrXq28 or a human-specific gRNA targeting chrXq22.1, in combination with 25*μ*M ML216 (Fig. S8, Materials and Methods).

Cells targeted with the chimpanzee-specific gRNA were sorted for the absence of mCherry, which marks the chimpanzee chrX, and increased intensity of GFP, to select for likely recombination events that result in two human alleles on the distal end of chrXq. Because we observed higher fluorescence intensity of the human chrX marker GFP in untreated cells during the G2/M cell cycle phase, we also used Hoechst DNA staining to sort specifically from G1 cells in the experiments with the chimpanzee-specific gRNA (Fig. S9, Materials and Methods). Similarly, cells targeted with the human-specific gRNA were sorted for the absence of GFP, which marks the human chrX, and increased intensity of mCherry. For the human-specific gRNA sorts, we also incorporated an additional marker by staining for a linked cell-surface protein, TSPAN6. Located on chrXq, *TSPAN6* has 1.4-fold higher *cis*-regulated expression from the chimpanzee allele compared to the human allele (adjusted *p* = 1.4×10^−4^ by Welch’s t-test); protein staining of TSPAN6 showed a similar difference (Fig. S9, Materials and Methods). Cells targeted with human-specific gRNA were thus sorted for absence of human-marker GFP and increased intensity of both chimpanzee-marker mCherry and of TSPAN6.

A total of 951 allo-tetraploid candidate colonies were grown from single cells after FACS. Additional genotyping confirmed that 172 colonies carried distal chrXq from a single species (Table S7). As expected, in 172/172 (100%) of these cases, the missing chrXq corresponded to the species targeted by the guide RNA (79/79 for the human gRNA, and 93/93 for the chimpanzee gRNA). To distinguish between deletion or recombination events, we determined the relative dosage of chrXq in these colonies by performing quantitative PCR (qPCR) assays on genomic DNA at chr6p, chrXp and chrXq (Table S2, SI Methods). We found that 171/172 (99.4%) colonies had lost the distal end of chrX of one species without altering the chrX dosage of the other species, as expected if targeting of the X chromosome had produced a species-specific deletion in these colonies.

We also identified one colony (0.6%) that had not only lost the distal end of the chimpanzee chrX but also doubled the dosage of the distal end of human chrX, consistent with a possible recombination event. Whole genome sequencing of this putative recombinant line confirmed that it was an inter-specific recombinant, with the first 140.1Mb of human chrX fused to the distal 27.6Mb of chimpanzee chrX (Xrec1 in Fig. 4B, Fig. S10). Sequence reads that span the precise junction between the human and chimpanzee sequences show that recombination did not occur in a region of large-scale homology between the two X chromosomes. Instead, a 4bp microhomology occurs directly at the junction site, suggesting that the recombination event was likely produced by microhomology-mediated end joining (MMEJ) and not by HR (29). No other human-chimpanzee recombinant chromosomes were found in the Xrec1 line when compared to an untreated control line.

We next leveraged the species-specific targeted lines as a panel of deletion lines for fine-mapping studies. We performed bulk RNA sequencing on 7 lines with partial chimpanzee chrX deletions, 8 lines with partial human chrX deletions, and 9 control lines without chrX deletions (Table S1). We identified the approximate breakpoint of each deletion by examining the ratio of reads that uniquely map to either the human or the chimpanzee genome along chrX (Fig. 4B, Fig. S11, Materials and Methods). In every case, mapped breakpoints were consistent with our results from genomic PCR and qPCR assays (Table S7). Three of the seven chimpanzee chrX deletions mapped within 1Mb of the chimpanzee-specific gRNA target site, and two of the eight human chrX deletions mapped within 1Mb of the human-specific gRNA target site (Fig. 4B). The remaining lines had a range of breakpoints that were up to 80Mb away from the species-specific guide targeting sites, and one cell line appeared to have lost the targeted X chromosome completely (Fig. 4B).

The gene *FMR1* is near the deletion breakpoints of three of the chimpanzee chrX deletion lines. We had identified in our bulk RNAseq analysis that *FMR1* is regulated in *cis*+*trans* between humans and chimpanzees with higher expression from the chimpanzee compared to human allele (Table S5). We similarly observed 1.5-fold higher expression from the chimpanzee *FMR1* allele compared to the human allele in our control lines without chrX deletions (adjusted *p* < 10^−12^ by the Wald test). As expected, all lines with human- or chimpanzee-specific deletion breakpoints proximal to *FMR1* had no expression from the deleted allele. One line (cXdel5) with a deletion breakpoint located 0.7-1Mb distal to *FMR1* showed significantly reduced *FMR1* expression (Fig. 4C). Two additional lines (cXdel6 and cXdel7) with deletion breakpoints 1.2-1.5Mb distal to *FMR1* showed normal *FMR1* expression. The reduced expression in cXdel5 is specific to the chimpanzee allele linked to the chimpanzee terminal deletion, suggesting that a *cis*-regulatory region that enhances *FMR1* expression maps to a 634kb interval (chrX:148656830-149290431, hg38) at least 0.7Mb downstream of *FMR1*.

If the X chromosome encodes *trans*-regulators of autosomal gene targets, partial deletions of either the human or chimpanzee X chromosome could result in significant gene expression changes for genes located on autosomes. Indeed, 42 autosomal genes showed significant changes in expression in the four deletion lines that removed regions on the chimpanzee X chromosome distal to breakpoints around 148Mb, and even more autosomal genes (147) showed significant changes in expression in the five cell lines that removed regions on the human X chromosome distal to breakpoints around 95Mb (Table S8). Interestingly, 7 of the genes altered by loss of distal chimpanzee chrX regions were not significantly different in the cell lines that had lost even larger regions of the human X chromosome. These genes also showed the expected signatures of a species-specific *trans*-effect when comparing gene expression levels among the different deletion lines (SI Methods). The autosomal genes included *MEGF10* and *TFPI2*, which were both classified as having a significant *trans*-component in our studies of intact diploid, auto-tetraploid, and allo-tetraploid cells (Fig. S12, Table S5). The magnitude of differential expression levels seen after species-specific removal of the distal X chromosome ranged from 60-80% of the overall expected *trans*-component. *Trans*-regulators encoded on the X chromosome may thus contribute to a fraction of the species-specific *trans*-expression differences observed in these autosomal genes. Extensions of this approach could be used to further localize both the responsible *trans*-factors on the X chromosome, as well as *trans*-factors on other chromosomes.

## Discussion

Understanding the molecular basis of human evolution is a grand and ambitious challenge in biological research. At the molecular level, researchers have cataloged the DNA sequence changes between humans and non-human primates (5) and identified many RNA expression differences between humans and chimpanzees across multiple tissues and developmental stages (13, 30, 31). However, it has been difficult to map the exact sequence changes that cause particular gene expression differences or other species-specific traits. Here, we have used intra- and inter-specific iPSC fusions to determine whether human-chimpanzee gene expression changes are controlled in *cis* or *trans*, and have developed genetic methods for further mapping both *cis*- and *trans*-regulatory changes to particular locations in the genome.

Regulatory changes appear to be a key driver of evolution in humans and other systems (32), and we and others have worked to determine the relative contribution of *cis*and *trans*-acting regulatory changes to gene expression differences between species. As in previous studies with human and chimpanzee iPSCs (13, 14), we found thousands of genes with species-specific expression differences. By comparing differential gene expression in single-species and cross-species fusions, we found that (1) both *cis-* and *trans*-regulatory changes are key contributors to human-chimpanzee differences, and (2) genes with *cis*-regulatory changes had, on average, more divergent expression than genes with *trans*-regulatory changes. Both of these findings are consistent with previous genome-wide studies of human-chimpanzee tetraploid cortical spheroids, human-chimpanzee tetraploid cranial neural crest cells, and inter-specific hybrids of mice, maize, *Arabidopsis*, and yeast (14, 15, 17, 33).

We also found that genes with *cis*-regulatory changes tended to influence fewer body parts than genes *conserved* between human and chimpanzee iPSCs. Furthermore, the number of body parts affected by a gene declines as the expression difference between humans and chimpanzees increases. *Cis*-regulatory changes are often thought to be favored in evolution because of their ability to avoid negative pleiotropy and restrict changes to particular tissues (32, 34). Our data suggest that genes that influence fewer biological processes are also more likely to evolve large expression differences as species diverge during evolution.

To facilitate further genetic mapping of human-chimpanzee differences, we examined multiple strategies to induce recombination events in allo-tetraploid iPSCs, including both genomewide and targeted approaches. The BLM inhibitor ML216 has been successfully used to induce inter-chromosomal recombination in other systems (11, 21, 23). In our experiments, ML216 treatments did not cause a measurable increase in intra-chromosomal exchange events scored by sister chromatid exchange assays, but may have stimulated a small ~12% increase on the number of human-chimpanzee recombinant molecules identified by haplotagging (Fig. 3). These differences could be non-significant, or might result from molecular differences between intra- and inter-specific recombination events; confounding effects of BrdU, which promotes DNA damage and differentiation (Fig. S13), in the sister chromatid exchange assay (35); or synergistic effects with CRISPR/Cas9 targeting, as previously reported for other mitotic recombination assays in human iPSCs (21). Further varying BLM activity by either pharmacological or genetic strategies (9, 10) could be tested for larger effects on the overall rate of recombination. Camptothecin and mitomycin C treatments are also promising candidates for further study, given their strong promotion of sister chromatid exchange events in allo-tetraploid iPSCs (Fig. 3B).

We also tested the ability of Cas9 and guide RNAs to stimulate inter-specific chromosome exchange events at particular locations in the genome. In contrast to prior work in yeast, *Drosophila*, and human iPSCs (12, 20, 21), we did not observe an enrichment in inter-specific recombination events at the site of targeting with or without ML216 by analyzing bulk populations with haplotagging. We also did not recover recombination events at the site of CRISPR targeting in the fluorescently-marked lines that we sorted to enrich for signatures of rare recombination events on the X chromosome. We did recover many lines that carried species-specific X chromosome deletions with breakpoints near the site of CRISPR targeting. We further recovered a single line carrying a confirmed human-chimpanzee recombinant X chromosome. However, the crossover point in the recombinant line was located tens of megabases away from the CRISPR targeting site and may be the result of a spontaneous or ML216-induced, rather than a CRISPR-induced, breakpoint on the X chromosome.

A variety of strategies may make it possible to increase the rate of homology-directed inter-specific recombination following CRISPR targeting. Following induction of doublestrand breaks by Cas9, homology-directed repair (HDR) pathways compete with several other pathways, including non-homologous end-joining (36). Studies in other systems have shown that the HDR pathway can be stimulated by expressing a plasmid with *RAD18*, a gene involved in the DNA damage response, or by treating cells with the small molecule RS-1 which increases the activity of the HDR-promoting protein RAD51 (37, 38). Conversely, the competing NHEJ pathway can be suppressed using the small molecule Scr7 to inhibit DNA Ligase IV, a key component of NHEJ (37). Studies in yeast show that tethering Cas9 to Spo11, a DSB-inducing protein with a key role in initiating meiotic recombination, can stimulate crossovers in naturally recombination-cold regions (39). These and other approaches can now be tested for their ability to stimulate targeted recombination between human and chimpanzee chromosomes in allo-tetraploid cells.

Like genetic mapping using recombinants, deletion mapping has also been used in many organisms to map phenotypes to specific genomic regions (40, 41). Our targeting and sorting strategies have already successfully produced a panel of deletion lines useful for further mapping of *cis*-regulatory elements and *trans*-acting factors on the X chromosome. The fraction of the genome removed by the induced chrX deletions is similar to the fraction of the genome removed by typical deficiency mapping chromosomes in *Drosophila* (0.2% of the genome deleted on average (41)). The staggered deletions on the chimpanzee X chromosome identify a previously unknown, long-distance, *cis*-regulatory region required for normal expression of the medically important *FMR1* gene. *FMR1* encodes an RNA-binding protein (FMRP) that is widely expressed in many tissues and modulates the translation of different sets of genes in specific contexts (42–44). Silencing of *FMR1* leads to Fragile X Syndrome, the most common form of intellectual disability in humans, but variation in *FMR1* expression can also result in other phenotypes including ataxia and primary ovarian insufficiency (45, 46). The 0.7Mb distance between *FMR1* and our newly-discovered downstream regulatory interval is large but not unprecedented. For instance, a key enhancer of Sonic Hedgehog (*SHH*) maps nearly 1Mb downstream of *SHH*, and mutations that disrupt this regulation lead to a variety of limb abnormalities in humans and other animals (47, 48). Interestingly, we found that *FMR1* is also expressed at different levels between human and chimpanzee iPSCs, a species difference that appears to come from a mixture of *cis*- and *trans*-acting regulatory changes. To determine whether the newly identified *cis*-regulatory interval underlies species differences in *FMR1* expression, simple extensions of our approach could be used to remove this downstream region specifically from the human X chromosome. Targeted deletion lines could also be used to look for species-specific translational changes that may provide insight into the effect of differential *FMR1* expression on human and chimpanzee biology.

Panels of chromosome deletion lines can also be used to map species-specific *trans*-regulators. *Trans*-effects appear to contribute to more than 50% of the gene expression differences identified between humans and chimpanzees in iPSCs (Fig. 2C), and are similarly pervasive in other systems (14, 15, 17, 33, 49). Our targeted X chromosome deletion lines suggest that human-chimpanzee differences in the autosomal genes *MEGF10* and *TFPI2* are controlled in part by species-specific *trans*-effects that map to the most distal ~8Mb of the X chromosome. One of the genes located in this distal X chromosome region is *MECP2*, which encodes a methyl DNA-binding protein that can activate or repress expression of target genes (50). Loss-of-function mutations in *MECP2* lead to Rett syndrome, a severe neurodevelopmental disorder. Intriguingly, prior research has identified both *MEGF10* and *TFPI2* as genes regulated by MeCP2 in human cells (51, 52). Further, both MEGF10 and MeCP2 have been linked to the pruning of neural synapses by astrocytes (53, 54), a cell type that has undergone changes in number, spatial organization, and function during human evolution (55, 56). Given that gene regulation in iPSCs cells has been shown to be similar to that in somatic tissues in some contexts (57), it is tempting to speculate that this potential *trans*-regulation might contribute to human-chimpanzee astrocyte differences or changes in neural processes and circuits pruned by astrocytes. Future experiments to selectively knock-out either the human or chimpanzee *MECP2* allele could test whether MeCP2 indeed regulates the species-specific expression of *MEGF10* and *TFPI2* in iPSCs, as well as potentially identify other species-specific *trans*-targets for this key transcriptional regulator.

We have focused most of our current studies on gene expression differences that are detectable in undifferentiated iPSCs. It is possible that tetraploidization will disrupt gene expression or limit the differentiation potential of auto- and allo-tetraploid iPSC lines. However, previous studies have shown that tetraploid mouse embryos can form most major organs, and rare humans with tetraploid karyotypes have been reported to survive for up to two years after birth (58, 59). In addition, our global RNA profiling experiments showed no large-scale gene expression disruptions between diploid and auto-tetraploid lines. We also find that diploid lines are more similar to their cognate auto-tetraploid lines than to other diploid lines of the same species. Thus, diploid and tetraploid iPSCs appear remarkably similar at the gene expression level. Future studies will be needed to determine if this similarity is maintained under a variety of differentiation conditions. Our initial experiments show that diploid, auto-tetraploid, and allo-tetraploid cells can all express characteristic gene markers of ectoderm, mesoderm, or endoderm under appropriate differentiation conditions (Fig. 1C, Table S3), and other tetraploid fusion lines have recently been differentiated into cortical spheroids or neural crest cells *in vitro* (14, 15). We caution that some of the lines in our own experiments showed incomplete endoderm differentiation, and previously-reported allo-tetraploid lines showed substantial expression of mesenchymal markers when incubated under conditions that stimulate cortical spheroid formation in diploids (14). *In vitro* differentiation protocols may thus need to be altered or optimized for tetraploid iPSCs to find conditions suitable for formation of particular cell types of interest.

Beyond mapping the *cis*- and *trans*-regulators of species-specific gene expression differences, we envision that allo-tetraploid iPSC lines will also be useful for mapping cellular and tissue differences between humans and chimpanzees. For example, many metabolic differences have evolved alongside major changes in diet between humans and chimpanzees (60). These changes are likely accompanied by cellular changes in enzyme levels and metabolite production that could be scored under appropriate *in vitro* conditions. In addition, neural progenitors in humans have been shown to have a longer prometaphase and longer metaphase compared to those in chimpanzees (61). These and other cellular traits can be assessed in culture and are compelling candidates for allo-tetraploid genetic mapping approaches. Recent advances in organoid technology also make it possible to study organ-level phenotypes that differ between humans and chimpanzees, including differences in organ size, connectivity, and cell type composition (31). Just as meiotic mapping panels have propelled our understanding of evolution in other organisms, further development of mapping methods in human-chimpanzee allo-tetraploids should provide powerful new genetic approaches for our quest to understand what makes us human.

## Materials and Methods

### Generation and maintenance of tetraploid iPSC lines

Human and chimpanzee diploid iPSC lines were labeled with diffusible fluorescent dyes and fused on an Eppendorf Multiporator at 4V AC for 80s, 16V DC for 20*μ*s, 6V post-AC for 95s (SI Methods). Tetraploid lines were confirmed by propidium iodide staining and karyotyping (Table S1). Diploid and tetraploid iPSC lines were routinely propagated feeder-free (SI Methods).

### Trilineage differentiation

Diploid and tetraploid iPSC lines were differentiated with the STEMdiff™ Trilineage Differentiation Kit according to the manufacturer’s instructions (STEMCELL Technologies, cat #05230). Differentiation was assessed using reverse transcription quantitative PCR (RT-qPCR) for pluripotency, ectoderm, mesoderm, and endoderm gene markers (Table S2, SI Methods).

### RNA sequencing analysis

Sequencing reads were aligned to a composite human-chimpanzee genome (hg38 and pt6), and the number of uniquely-mapped reads that overlap each gene was determined using a curated exon annotation (SI Methods). Differential gene expression analysis between diploids and auto-tetraploid iPSCs was performed with DESeq2 (62), and genes with an adjusted *p* < 0.05 and at least a 2-fold change in expression were called as significant (SI Methods).

Differential expression (DE) between single-species iPSCs, allelespecific expression (ASE) in allo-tetraploids, and regulatory type classifications were carried out as a combination of previously described methods (33, 49). Genes were classified as *cis, trans, cis+trans, cis-trans, compensatory, conserved*, or *ambiguous* based on different combinations of significant DE, significant ASE, significant *log*_2_(*FC*) difference between DE and ASE (“*trans*-effects”), and direction of *cis*-contribution and *trans*-contribution to the DE *log*_2_(*FC*) (SI Methods).

### Sister chromatid exchange (SCE) assay

Camptothecin (Sigma Aldrich, cat #C9911-100MG), ML216 (Cayman Chemical, cat #15186), and mitomycin C (Sigma Aldrich, cat #M4287-2MG) were applied to iPSCs with 10*μ*M BrdU (SI Methods). The sister chromatid exchange (SCE) assay was then performed as previously described (24).

### Haplotagging

Haplotagging was performed as previously described (27). Reads were aligned to a composite human-chimpanzee genome (hg38 and pt6) and assigned to their molecule of origin by barcode. Variants between hg38 and pt6 were identified for each read and filtered by multiple criteria (SI Methods). Molecules were scored as recombinant if they contained 1 inter-specific event and at least 2 supporting variants per species.

### FACS of fluorescently-marked allo-tetraploid lines

Using two rounds of homologous recombination, we inserted an EF1a-EGFP-IRES-PuroR cassette at human chrXq28 and an EF1a-mCherry-IRES-NeoR cassette at chimpanzee chrXq28 in allo-tetraploid iPSCs (Fig. S7, SI Methods). Double-marked iPSCs were treated with 25*μ*M ML216 starting 12 hours before nucleofection of CRISPR/Cas9 and gRNA and continuing until 48 hours post-nucleofection. We then employed multiple sorting strategies to enrich for chrX recombination or deletion events (Fig. S9, SI Methods).

### DNA sequencing analysis to identify chrX recombination site

DNA sequencing reads from the recombinant allo-tetraploid cell line, H1C1a-X1-Xrec1, and a control allo-tetraploid line, H1C1a-X1-S, were aligned to a composite human-chimpanzee (hg38-pt6) reference genome (SI Methods). The read counts in H1C1a-X1-Xrec1 were normalized to read counts in H1C1a-X1-S (Fig. S10).

## Data Availability

Data supporting the findings of this study are included in the main text and SI Appendix or deposited in publicly available databases. RNA sequencing data generated in this study are available at GEO (pending), and the DNA sequence containing the recombination site for H1C1a-X1-Xrec1 is available at Genbank (pending). Additional materials will be made available upon request.

## ACKNOWLEDGMENTS

We thank Yoav Gilad for human and chimpanzee iPSC lines, and the Stanford Shared FACS Facility, WiCell Research Institute, Kyle Loh, Alyssa Benjamin, and members of the Kingsley and Chan labs for useful discussions. This work was supported in part by pre-doctoral fellowships from the National Science Foundation (J.H.T.S., R.L.G., G.A.R.K.), by Stanford Graduate Fellowships (J.H.T.S., R.L.G., G.A.R.K.), by a fellowship from the Center for Computational, Evolutionary and Human Genomics at Stanford (J.H.T.S.), by National Institutes of Health grant T32GM007790 (V.C.B.), and by European Research Council Grant “HybridMiX” 639096 (Y.F.C). Y.F.C. is supported by the Max Planck Society, and D.M.K. is an investigator of the Howard Hughes Medical Institute.

## Supplementary Information

### Supporting Information Text

#### Supplemental Materials and Methods

##### Cell culture maintenance

The induced pluripotent stem cell lines H23555 (H1), H20961 (H2), C3649 (C1), and C8861 (C2) were provided by the Gilad laboratory (1). Cultures were tested for and maintained mycoplasma free. Diploid and tetraploid lines were routinely propagated feeder-free in mTeSR1 or mTeSR Plus media (STEMCELL Technologies, cat #85850 and cat #100-0276) on cell culture plastics coated with Geltrex basement membrane matrix (Gibco, cat #A1413302). When confluent, cells were passaged using Accutase (Millipore, cat #SCR005) with 1*μ*M thiazovivin (Tocris, cat #3845) or using 0.5mM EDTA as previously described (2, 3). Cells were imaged on the EVOS FL microscope (Thermo Fisher) at 4X magnification, unless otherwise noted.

##### Generation of tetraploid iPSC lines

One diploid iPSC line was labeled with CellTracker Green CMFDA Dye (1:667) (Thermo Fisher, cat #C7025) and the other diploid iPSC line was labeled with CellTracker Blue CMAC dye (1:500) (Thermo Fisher, cat #C2110) or CellTracker Red CMTPX Dye (1:1000) (Thermo Fisher, cat #C34552) per the manufacturer’s instructions. Cells were washed multiple times to remove excess dye and allowed to recover after labeling in mTeSR1 + 1*μ*M thiazovivin for at least 1 hour prior to fusion. 7*x*10^5^ cells from each line were combined, washed twice in 1ml fusion buffer, resuspended in 350*μ*l fusion buffer, and fused in the helix fusion chamber of an Eppendorf Multiporator at 4V AC for 80s, 16V DC for 20*μ*s, 6V post-AC for 95s. After 10 minutes at room temperature, 1ml of mTeSR1 + 1*μ*M thiazovivin was added to the helix fusion chamber. Fusion buffer was hypoosmolar electrofusion buffer (Eppendorf, cat #940002150) diluted in water (normally, 60% hypoosmolar electrofusion buffer and 40% water).

Tetraploid clones were then selected in one of two ways. In the first method, 250-350*μ*l of the resulting suspension after fusion was immediately plated in a 10-cm plate. Double-labeled cells were screened the following day under a fluorescent microscope and marked on the plate. The diffusible dyes were only visible for 2 days after fusion. Surrounding diploid cells were removed on each subsequent day by manual scraping. When the originally identified double-labeled cells grew into colonies, they were picked into a 96-well plate and screened as described below.

In the second method, 7-15 fusions were performed on the same day using the helix fusion chamber as described above and collected in a large volume of mTeSR1 + 1*μ*M thiazovivin for fluorescence-activated cell sorting (FACS). For increased viability in the second method, we used only CellTracker Green CMFDA and CellTracker Blue CMAC dyes. Cells were gently pipetted every hour to avoid CellTracker dye diffusion and undesired labeling of nearby non-fused cells. Prior to FACS, cells were resuspended in 500*μ*l FACS buffer (0.5%BSA fraction V, 5mM EDTA, 1% Penicillin/Streptavidin, and 1*μ*M thiazovivin in 1X PBS) and strained through a 30*μ*m filter. Double-labeled cells were collected into 1 well of a 96-well plate by FACS. After 3 days, cells were collected and seeded at 1-5*x*10^3^ cells in 10-cm plates. Individual colonies were then picked into 96-well plates.

To identify tetraploid colonies, we performed propidium iodide staining (Invitrogen, cat #P3566) on fixed cells to examine ploidy via FACS analysis. Briefly, cells were fixed in 80% EtOH overnight at 4°C. The next day, cells were washed twice in 1X PBS and stained at 37°C for 10 minutes in 20*μ*g/ml RNase A, 40*μ*g/ml of propidium iodide, and 0.1% Triton X-100 in 1X PBS. Cells with both 4N and 8N DNA content by FACS, suggesting that they contain tetraploid cells, were expanded. Expanded colonies were further screened for DNA content by karyotyping as described previously (4). Colonies that contained only tetraploid cells were maintained as stocks. G-banded karyotyping was also performed by WiCell (Table S1).

##### Trilineage differentiation

Diploid and tetraploid iPSC lines were seeded in 12-well plates and differentiated with the STEMdiff™Trilineage Differentiation Kit according to the manufacturer’s instructions (STEMCELL Technologies, cat #05230). Additionally, untreated cells were collected two days after seeding. Three replicate wells of cells per cell line were collected per condition. Differentiation was assessed using reverse transcription quantitative PCR (RT-qPCR) for pluripotency, ectoderm, mesoderm, and endoderm gene markers (Quantitative PCR SI Methods).

##### Quantitative PCR (qPCR)

RT-qPCR was used to assess differentiation potential for trilineage differentiation samples, and qPCR was used to study DNA marker dosage in chrX targeted cell lines. All reactions were performed using Brilliant II SYBR Green Low ROX qPCR Master Mix (Agilent, cat #600830) on a QuantStudio 5 Real-Time PCR System (Thermo Fisher).

For trilineage differentiation, three replicate wells of cells per condition were collected in Trizol and applied to the Direct-Zol RNA Miniprep kit (Zymo Research, cat #R2051) for RNA extraction per the manufacturer’s instructions. cDNA was synthesized from RNA with the SuperScript^TM^VILO^TM^ cDNA Synthesis Kit (Thermo Fisher, cat #11754250). To assess differentiation potential of trilineage differentiation samples, qPCR was performed in triplicate on all samples with two marker genes each for pluripotency (*NANOG, DNMT3B*), ectoderm (*PAX6, RAX*), mesoderm (*TBXT, HAND1*), endoderm (*FOXA2, SOX17*), and housekeeping (*GAPDH, YWHAZ*) (5–11). All primer pairs span a large intron, have efficiencies between 90-110%, bind identical sequences in humans and chimpanzees, produce PCR products of identical length in both species, and were chosen from the literature or designed in-house (Table S2). For each sample, the quantity of each marker gene was calculated by comparing to a standard curve of pooled samples. This quantity was normalized by dividing by the geometric mean of the quantities of the two housekeeping genes (*GAPDH*, *YWHAZ*) in the same sample and then divided by the normalized quantity of the marker gene in undifferentiated iPSCs from the H2 human diploid line. Two-tailed Student’s t-tests were used to determine statistically significant differences in marker gene expression between differentiated and undifferentiated iPSCs at 5% Benjamini-Hochberg FDR.

For determination of chrXq dosage relative to other chromosomes, cells were harvested from 96-well plates using Accutase (Millipore, cat #SCR005), and DNA was extracted using the DNeasy 96 Blood & Tissue Kit (Qiagen). Reactions were performed either in duplicate or in triplicate with primers for chromosomes 6p, Xp and Xq (Table S2).

##### Library preparation for RNA sequencing

Samples were flash frozen and stored at −80°C as a pellet. RNA extraction, library preparation, and sequencing for the chrX deletion samples were performed by Genewiz. All other RNA sequencing samples were prepared in-house before sequencing on the Illumina HiSeq 4000 with Novogene. Briefly, samples were resuspended in Trizol and directly applied to the Direct-Zol RNA Miniprep kit (Zymo Research, cat #R2051) for RNA extraction per the manufacturer’s instructions. Technical replicates for each line were collected from thaws of different frozen vials. Only samples with *RIN* > 9 were used for RNA sequencing.

1*μ*g of RNA was used for library preparation. RNA sequencing libraries were prepared with the TruSeq Stranded mRNA Library Prep (Illumina, cat #20020595) using the IDT for Illumina – TruSeq RNA UD Indexes (96 Indexes, 96 Samples) (Illumina, cat #20022371) according to the manufacturer’s instructions with one modification. Prior to PCR amplification, 10% of a subset of samples were run under the recommended PCR conditions with SybrGreen on the QuantStudio 5 Real-Time PCR System. We identified the number of PCR cycles required to reach the crossing point by qPCR and used that number of cycles for PCR amplification on the entire set of samples. We ran 8 PCR cycles. Libraries were pooled and sequenced to around 10 million reads per sample for the diploid and auto-tetraploid lines and around 20 million reads per sample for the allo-tetraploid lines. Five independently-derived C1C1 auto-tetraploid lines, two independently-derived H1H1 auto-tetraploid lines (two technical replicates each), twelve independently-derived H1C1 allo-tetraploid lines, and ten independently-derived H2C2 allo-tetraploid lines were sequenced. Three technical replicates were sequenced for each diploid line.

##### Alignment of RNA sequencing to composite human-chimpanzee genome

Sequencing reads were trimmed for adapter sequences using cutadapt v1.8.1 (12), and read quality was confirmed using fastqc v0.11.9 (13). Reads were aligned using STAR v2.7.1a with two-pass mapping (14). Samples were mapped to a composite human-chimpanzee genome (hg38 and pt6). The number of uniquely-mapped reads (*MAPQ* = 255) that overlap each gene was counted using featureCounts from the subread v1.6.0 package (15).

To generate the gene annotations used in featureCounts, GRCh38.94 human exon annotations from Ensembl (16) were mapped from hg38 to pt6 using pslMap (17). After removing mappings where the number of bases that map is less than half of the query exon size, we retained only exons that uniquely mapped from humans to chimpanzees. We then removed genes for which exons map to opposite DNA strands, different scaffolded chromosomes, or where consecutive exons map more than 800kb apart. We further filtered out exons where more than 10% of reads from diploid or auto-tetraploid lines map to the incorrect species when mapped to the composite genome. This resulted in 48,735 annotated genes that contain at least 1 exon (byexon-gene). We also used a second set of annotations. We identified SNPs that differed between the human and chimpanzee cell lines using the GATK RNA variant pipeline (18, 19) and assigned SNPs to genes annotated in humans. We also filtered out SNPs where more than 10% of reads from diploid or auto-tetraploid lines map to the incorrect species when mapped to the composite genome. This resulted in 14,333 annotated genes with at least 1 SNP (bysnp-gene). Read counts for the byexon-gene annotation were also adjusted for feature length to account for differences between feature length in the human and chimpanzee genomes. Results were very similar across both annotations, and results are reported for the byexon-gene annotation in the current study.

##### Gene expression analysis of diploids and auto-tetraploid iPSC lines

After sequencing reads were aligned to the composite human-chimpanzee genome as described above, differential gene expression analysis was performed with DESeq2 (20) using default parameters. We called genes as significant if they had an adjusted *p* < 0.05 after Benjamini-Hochberg FDR correction and at least a 2-fold change in expression.

##### Differential gene expression, allele-specific gene expression, and *cis/trans* analysis between humans and chimpanzees in diploid, auto-tetraploid, and allo-tetraploid iPSC lines

Differential expression (DE) between single-species iPSCs, allele-specific expression (ASE) in allo-tetraploids, and regulatory type classifications were carried out as a combination of previously described methods (21, 22).

After RNA sequencing reads were aligned to the composite human-chimpanzee genome as described above, reads mapping to genes on human chromosome 18 (and the orthologous chimpanzee genes) were removed. Next, each sample was downsampled to 9,711,244 reads (for DE) or 11,490,119 reads (for ASE), and genes with fewer than 10 reads assigned to both the human and the chimpanzee orthologs were excluded. For each gene, *log*_2_(*FC*) was calculated between each human-only and each chimpanzee-only sample (for DE) or between the human allele and the chimp allele in each allo-tetraploid cell line (for ASE). Genes with significantly different *log*_2_(*FC*) between human and chimpanzee were determined to be DE or ASE. Each gene was tested for significant “*trans*-effects” by testing for a significant *log_2_(FC*) difference between single-species iPSCs and allo-tetraploid iPSCs. Significance for all *log_2_(FC*) differences was determined by Welch’s t-test at 5% Benjamini-Hochberg FDR. Importantly, only half of the allo-tetraploid samples were used to determine whether a gene is significantly ASE, and the other half were used to determine significant “*trans*-effects” since this has been reported to reduce false classification as *compensatory* (23).

Finally, the *cis*-contribution (*C*) and *trans*-contribution (*T*) to the observed DE *log*_2_(*FC*) (*D*) was calculated for each gene. Specifically, the *cis*-contribution (*C*) was equal to the ASE *log*_2_(*FC*), and the *trans*-contribution was calculated as *T* = *D* – *C*.

Genes were classified by regulatory type based on the following criteria:

***cis***: significant DE, significant ASE, no significant “*trans*-effects,”*cis*-contribution and *trans*-contribution to DE *log*_2_(*FC*) in the same direction
***trans***: significant DE, not significant ASE, significant “*trans*-effects”
***cis+trans***: significant DE, significant ASE, significant “*trans*-effects,” *cis*-contribution and *trans*-contribution to DE *log*_2_(*FC*) in the same direction
***cis-trans***: significant DE, significant ASE, significant “*trans*-effects,” *cis*-contribution and *trans*-contribution to DE *log*_2_(*FC*) in opposite directions
***compensatory***: not significant DE, significant ASE, significant “*trans*-effects”
***conserved***: not significant DE, not significant ASE, no significant “*trans*-effects”
***ambiguous***: all other patterns

All results reported in this paper used the “by-exon-gene” annotation as described in the “Alignment of RNA sequencing to composite human-chimpanzee genome” section above (except for *TSPAN6*, which was not included in the “byexon-gene” annotation and was assessed using the “bysnp-gene” annotation).

##### Gene ontology enrichments

Significant gene ontology enrichments (adjusted *p* < 0.05 after Benjamini-Hochberg FDR correction) were determined using the R package clusterProfiler’s enrichGO function (24) for the annotation data sets “Biological Process,”“Molecular Function,” and “Cellular Component.” The set of analyzed genes was used as the background reference list.

##### Gene expression analysis of X chromosome deletion lines

RNA sequencing reads were aligned to the composite human-chimpanzee genome as described above. To identify the approximate location of X chromosome deletions, we computed the ratio of human read counts to chimpanzee read counts for each deletion line normalized to control (non-deletion) lines. A count of 1 was added to any sample with allele counts of zero, and ratios were calculated for genes with more than 10 counts on average and where at least half of the samples had at least 5 counts. Approximate deletion breakpoints were then determined by visual inspection.

To identify autosomal genes whose expression may be affected by *trans*-regulators on the X chromosome, we carried out differential gene expression analysis of control and human and chimpanzee chrX targeted deletion lines using DESeq2 (20) with the Wald test at 5% Benjamini-Hochberg FDR. *Trans*-regulated candidates were identified by the following five criteria: (1) Genes on autosomes that showed significant expression changes when comparing the four lines with deletion breakpoints of the chimpanzee chrX around 148Mb (cXdel4-cXdel7) to the nine control lines that lack deletions; (2) Genes on autosomes that did not show significant expression changes when comparing the five lines with deletion breakpoints of the human chrX around 95Mb (hXdel3-hXdel7) to the nine control lines that lack deletions; (3) Genes that met the first two criteria whose expression level was also significantly different in comparisons between the chimpanzee (cXdel4-cXdel7) and human (hXdel3-hXdel7) terminal deletions; (4) Genes that also showed the same direction of change in cell lines carrying shorter (cXdel4-cXdel7) and larger chimpanzee chrX deletions (cXdel1-cXdel3) compared to control lines; and (5) Genes where the hXdel8 line which has a human deletion breakpoint near the distal chimpanzee chrX deletion lines maintained expression within the range of the control lines.

##### Sister chromatid exchange (SCE) assay

Cells were passaged the day before testing. For camptothecin (Sigma Aldrich, cat #C9911-100MG), camptothecin and 10*μ*M BrdU were applied to cells for 1 hour before being replaced with fresh media containing 10*μ*M BrdU overnight. For ML216 (Cayman Chemical, cat #15186) and mitomycin C (Sigma Aldrich, cat #M4287-2MG), cells were incubated with the small molecule and 10*μ*M BrdU for 24-48 hours with a media change every 24 hours. Cells were then moved to fresh media containing 10*μ*M BrdU and 0.1*μ*g/ml colcemid for 4 hours and subsequently collected for sister chromatid exchange (SCE) assay as previously described (25). Cells were alternatively first collected into a 1.5ml tube before adding new media containing 10*μ*M BrdU and 0.1*μ*g/ml colcemid, with no obvious change in results. Multiple metaphase spreads were imaged at 100X, and recombination events were counted using the ImageJ Cell Counter function. *P*-values were calculated using the 1-tailed Student’s t-test.

##### Haplotagging

Haplotagging was performed as previously described (26). Briefly, genomic DNA from each sample was mixed with individually barcoded magnetic beads containing bead-immobilized active Tn5 transposase for tagmentation with up to 21 million barcode diversity. Tagged DNA was then PCR amplified, size selected, and sequenced on a NovaSeq 6000 instrument (Illumina).

Reads were aligned to a composite human-chimpanzee genome (hg38 and pt6) using EMA, a barcode-first variant of the bwa aligner (27). For the analysis, we focused on regions that reciprocally and uniquely mapped between the two species assemblies, with the mapping based on the hg38 to pt6 chain files from the UCSC Genome Browser (28) and pslMap (17). 500bp orthologous regions with greater than 2-fold difference in read coverage were excluded from further analysis. Each read was also assigned to a molecule based on its barcode (retained as the BX beadTag). For each read, we identified variants between hg38 and pt6 (SNPs and indels). The variant annotation file was generated by first parsing the maf file between hg38 and pt6 from the UCSC Genome Browser (28). We also included variants identified by running the GATK variant pipeline (18, 19) on reads that map uniquely to either hg38 or pt6 and where all reads assigned to a given barcode map to only one species. If no variants in our resulting annotation file were identified in a read but the read uniquely mapped to either hg38 or pt6, the read itself was considered as a variant.

Along each molecule, we coded the species assignment (e.g *H-H-H-H-H-C-C-C* where *H* is a human variant and *C* is a chimpanzee variant). We then applied the following strict filters to identify a high-confidence set of recombinant molecules: (1) Identified SNPs must have a phred quality score of at least 30; (2) Given the low rate of mitotic recombination, multiple “switch” events (e.g. *H-H-H-C-H-H-H*) are likely artifacts and such variants were removed; (3) We also excluded possible mapping artifacts where particular variants were found at the boundary of multiple recombination events; (4) Variants contained in 500bp regions with greater than 2-fold difference in the directionality of switch events (e.g. switch events were predominantly *H*→*C* instead of *C*→*H*) were removed; (5) We included only paired recombinant molecules that could be “reciprocal events” to further account for biases in the directionality of switch events; (6) We excluded any variants that are in ENCODE blacklist regions (29); (7) All recombinant molecules must contain only 1 switch event and at least 2 supporting variants per species.

To assess the effect of CRISPR targeting to specific loci, we examined the recombination rate in the 250kb interval surrounding the target loci with and without filters, with no difference in the relative enrichment at the target loci. Data visualizations were generated with ggplot2 (30) and karyoploteR (31). In Fig. S6, samples were plotted with their experimental batch due to differences in read and molecular coverage.

##### Generation of fluorescently-tagged allo-tetraploid lines

We cloned two plasmids, one with homology arms (chrX:153,850,316-153,851,493, hg38) flanking a EF1a-EGFP-IRES-PuroR cassette to target human chrX and the second with homology arms (chrX:149,205,726-149,208,867, pt6) flanking a EF1a-mCherry-IRES-NeoR cassette to target chimpanzee chrX (Fig. S7), into the pMAXGFP plasmid backbone (Lonza). Guide RNAs (gRNAs) were designed to linearize the plasmid containing the insertion cassette and cut the target insertion site (HR_X_gRNA_1 and HR_X_gRNA_2 in Table S2). gRNAs were then *in vitro* transcribed as described above.

2.5*μ*l of 40*μ*M Cas9-NLS purified protein (QB3, UC Berkeley) was mixed with 2.5*μ*g each of both gRNAs for 10 minutes at room temperature. This complex and 1.875*μ*g of the plasmid targeting human chrX were nucleofected into 3*x*10^6^ cells using the Nucleofection Stem Cell Kit 2 (Lonza, cat #VPH-5022) and program A-33 on the Nucleofector 2b Device (Lonza). Immediately after nucleofection, 1ml of pre-warmed media (mTeSR1 + 1*μ*M thiazovivin) was added to the reaction. The reaction was allowed to recover for 20 minutes at room temperature, and 5 separate reactions were pooled and plated on one 10-cm plate. We also nucleofected pMAXGFP separately as a positive control for nucleofection efficiency.

After cells recovered and expanded (~5 days post-nucleofection), cells with insertion events were selected by multiple days of puromycin treatment. We examined selection efficacy via fluorescence under an EVOS FL microscope. After multi-day selection, we picked colonies into 96-well plates. When colonies reached confluency, they were split and screened for proper insertion events by PCR, using primer pairs where one primer targets nearby genomic DNA and a second primer targets the insertion construct. We verified target-site insertion events using primer sets at both the 5’ and 3’ ends and separately with species-specific primers (Table S2). We confirmed the insertion sequence via PCR followed by Sanger sequencing with primers chrX-F2 and chrX-R2 (Table S2). To confirm that the insertion was inserted into the target locus and nowhere else in the genome, we expanded promising colonies for Southern blot analysis. Colonies verified by both PCR and Southern blot were then subject to a second round of nucleofection to insert the mCherry cassette into chimpanzee chrX. Double-marked colonies were selected for using Geneticin (Thermo Fisher, cat #10131035) and puromycin (Sigma-Aldrich cat #P8833). Double-marked lines were confirmed by PCR, Southern blot, and visual inspection of fluorescence.

##### CRISPR/Cas9 treatment of iPSC lines

Guide RNAs were designed (Table S2) and *in vitro* transcribed (IVT) as previously described (32). Briefly, CRISPR IVT target oligos containing the gRNA and the CRISPR IVT scaffold oligo (HPLC-purified) were synthesized by Integrated DNA Technologies. 40 cycles of PCR were performed between the CRISPR IVT scaffold oligo using Phusion DNA polymerase (Thermo Scientific, cat #F530L) and the CRISPR IVT target oligo, and the PCR product was purified using the QIAquick PCR Purification Kit (Qiagen, cat #28104). The PCR product was then *in vitro* transcribed using the MEGAscript T7 transcription kit (Thermo Fisher, cat #AM1334) for 16 hours at 37°C. The reaction was treated with DNase, and transcribed gRNA was extracted with phenol/chloroform and precipitated with isopropanol. Transcribed gRNA was resuspended to approximately 2*μ*g/ul.

To select the highest efficiency guides, we tested performance in 96-well plate format. For each guide, 1*μ*l of 40*μ*M Cas9-NLS purified protein (QB3, UC Berkeley) was complexed with 2*μ*g of gRNA for 10 minutes at room temperature. We performed nucleofection of 2*x*10^5^ cells per reaction using the P3 Primary Cell 96-well Nucleofector Kit (Lonza, cat #V4SP-3096) on the Amaxa 96-well Shuttle Device (Lonza) with program CA-137. Two days post-nucleofection, cells were collected for DNA extraction using phenol/chloroform. We used primers bracketing the target cut site (Table S2) and Sanger sequenced the products for analysis using TIDE (33) to determine guide efficiency. For a subset of guides, we further confirmed cutting events by cloning the gel-extracted PCR product into the TOPO TA vector (Life Technologies, cat #450641) and performing colony PCR followed by Sanger sequencing to identify lesions at the target cut site.

For targeted recombination, we nucleofected cells with CRISPR/Cas9 and gRNA using the same nucleofection conditions as described above in the “Generation of fluorescently-tagged allo-tetraploid lines” section. For CRISPR+ML216 conditions, we treated cells with 25*μ*M ML216 starting 12 hours before nucleofection, as previously described (34). After recovering for 1 hour, nucleofected cells were plated directly into media with ML216, and ML216 media was replaced again after 24 hours. At 48 hours post-nucleofection, cells were collected for FACS or haplotagging experiments.

##### Fluorescence activated cell sorting (FACS)

Allo-tetraploid cells with fluorescently-marked chrX were subjected to CRISPR+ ML216 treatment as described above.

For cells treated with chimpanzee-specific gRNA, we selected for loss of mCherry (from the chimpanzee chrX insertion) and possible duplication of GFP (from the human chrX insertion) by sorting for mCherry-negative, high-intensity-GFP cells. To eliminate cells with high GFP due to duplicated DNA content at the G2/M phase of the cell cycle, we stained cells with Hoechst 33342 (Thermo Fisher, cat #62249) for 30 minutes at 37°C to sort only from cells in the G1 cell cycle phase.

For cells treated with human-specific gRNA, we selected for loss of GFP (from the human chrX insertion) and possible duplication of mCherry (from the chimpanzee chrX insertion) by sorting for GFP-negative, high-intensity-mCherry cells. As a second marker of two copies of chimpanzee chrX downstream of the gRNA cutsite, we chose a cell-surface protein, TSPAN6, that has *cis*-regulatory changes with 1.4-fold higher expression in chimpanzee relative to human (adjusted *p* = 1.4<10^−4^ by Welch’s t-test after Benjamini-Hochberg FDR correction). Because this human-chimpanzee gene expression difference can also be observed at the level of protein expression by antibody staining, sorting for high TSPAN6 protein acted as a second marker to potentially sort for two copies of chimpanzee chrX downstream of the gRNA cutsite (Fig. S9).

Treated cells were stained with either Hoechst 33342 at 10*μ*g/mL for 30 minutes at 37°C, or with TSPAN6 primary antibody (1:10; LS Bio, cat #LS-C160272-400) for 1 hour at 4°C followed by 30 minutes at 4°C with a goat anti-rabbit secondary antibody conjugated with APC fluorophore (1:500; Thermo Fisher, cat #A-10931). Cells were sorted single-cell into 96-well plates on a BD Influx cell sorter at the Stanford Shared FACS Facility. Representative sorting gating schemes are shown in Fig. S9.

##### DNA sequencing analysis to identify chrX recombination site

DNA from the recombinant allo-tetraploid cell line, H1C1a-X1-Xrec1, and a control allo-tetraploid line, H1C1a-X1-S, (Table S1) were extracted and sequenced to 30X coverage with 150bp paired-end reads by GeneWiz. Illumina adapters were removed using Picard Tools (http://broadinstitute.github.io/picard/), and reads were then aligned to a composite human-chimpanzee (hg38-pt6) reference genome using BWA-MEM with the -M flag (35). Duplicate reads were marked with Picard Tools and removed using samtools (36). We filtered out reads with *MAPQ* < 30 and reads that did not lift over between hg38 and pt6 using pslMap (17). To identify the likely recombination site, we normalized the read counts in H1C1a-X1-Xrec1 to read counts in the control H1C1a-X1-S over 10kb sliding windows to account for any sequencing or mapping bias and visualized this ratio along human chrX coordinates (Fig. S10). Inspection of reads at the likely recombination site revealed the exact junction site as a 4bp microhomology (CACC) found at both human chrX:140133478-140133481 (hg38) and chimpanzee chrX:124020937-124020940 (pt6).

---

**Fig. S1.**
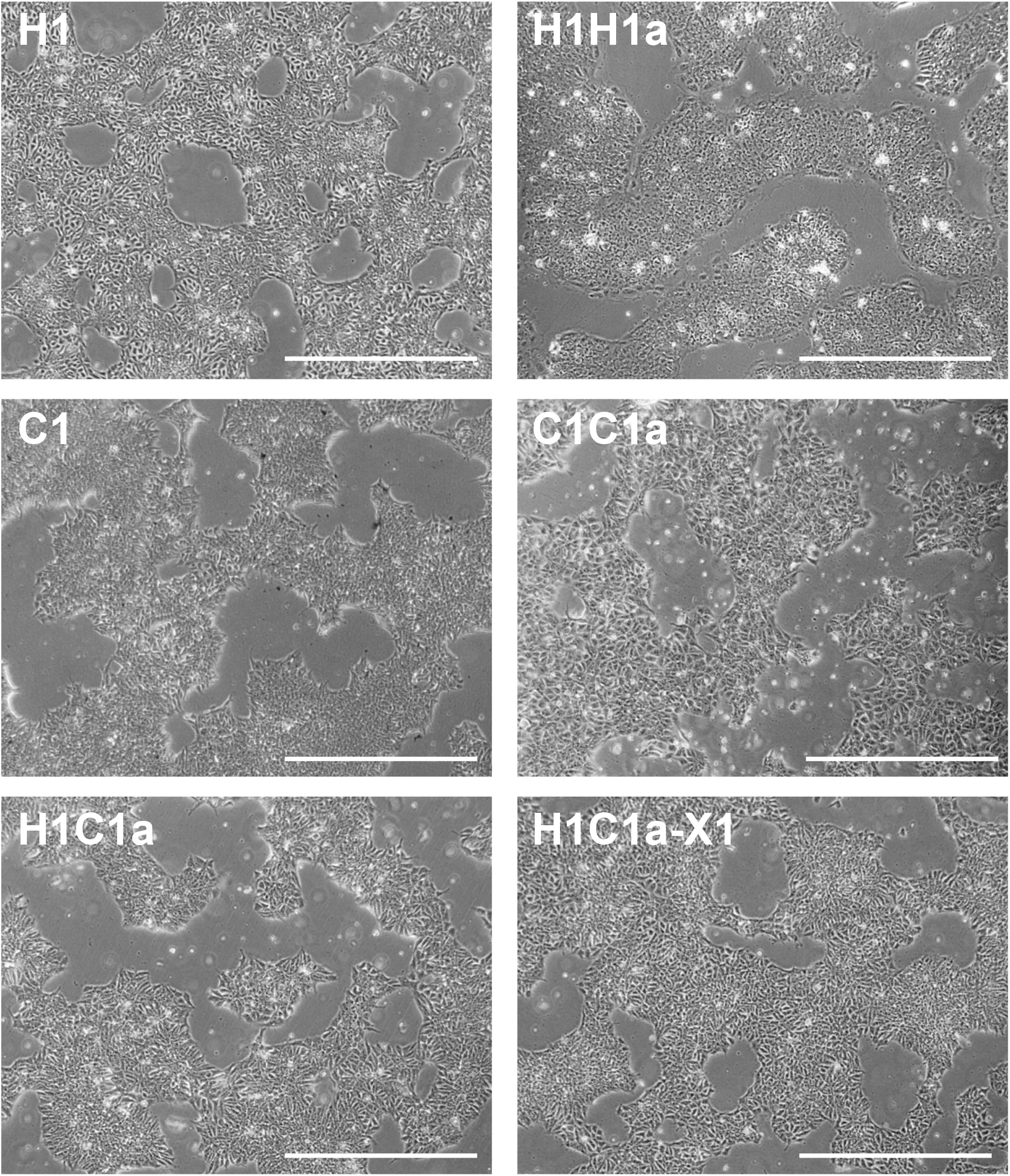
Morphologies of auto- and allo-tetraploid iPSC lines are similar to those of diploid iPSC lines. Representative brightfield images of human diploid (H1), human auto-tetraploid (H1H1a), chimpanzee diploid (C1), chimpanzee auto-tetraploid (C1C1a), allo-tetraploid (H1C1a), and chrX-marked allo-tetraploid (H1C1a-X1) lines are shown. Scale bars are 1mm.

**Fig. S2.**
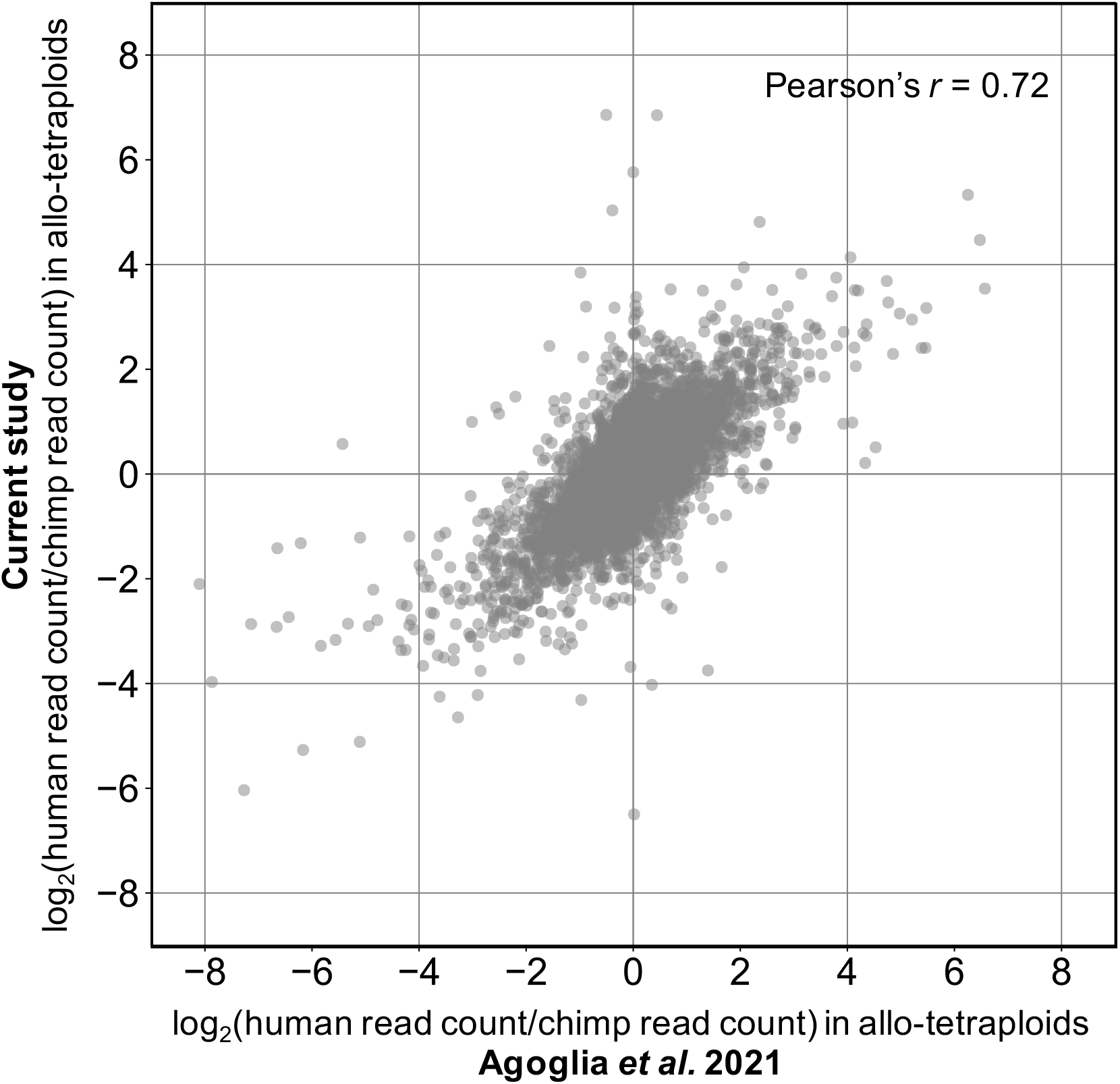
Allele-specific expression in human-chimpanzee allo-tetraploid iPSCs is reproducible across studies. Allele-specific expression (ASE) *log*_2_(*FC*) values from RNAseq generated by Agoglia *et al*. 2021 (37) (x-axis) and ASE *log*_2_(*FC*) values from the RNAseq data reported in this study (y-axis) are highly concordant (Pearson’s *r* = 0.72). Allo-tetraploid cells were derived from independent human-chimpanzee iPSC fusion events in the two studies, and different pipelines were used for mapping reads, assigning reads to the human or chimpanzee version of a gene, and calling genes with significant ASE. ASE differences in human-chimpanzee allo-tetraploid iPSCs are thus highly reproducible and robust to different analysis methods.

**Fig. S3.**
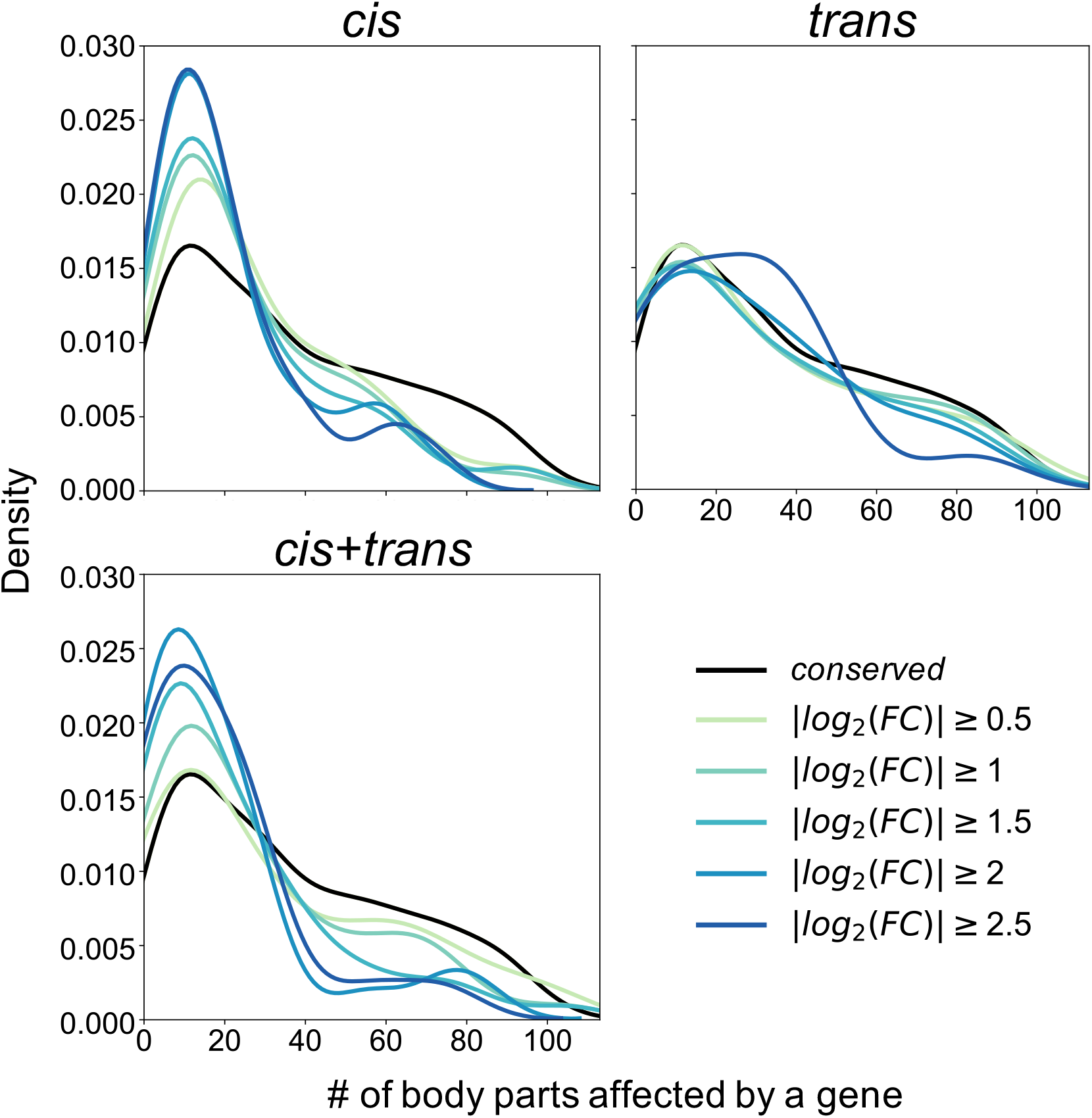
Genes with increasingly divergent expression between human and chimpanzee iPSCs influence fewer body parts for *cis* and *cis+trans* regulatory types. Density plots (smoothed histograms) showing the distribution of body parts influenced by genes (according to the Gene ORGANizer database (38)) with human-chimpanzee expression differences due to *cis* (upper left), *trans* (upper right), and *cis+trans* (lower left) regulatory changes at increasing |*log*_2_(*FC*)| cutoffs. The *cis-trans* category is not included because only 5 genes have |*log*_2_(*FC*)≥1|. For genes classified as *cis* and *cis+trans*, the median number of body parts influenced decreases with higher |*log*_2_(*FC*)| cutoffs (22, 18, 17, 15, 15 body parts and 22.5, 18, 16, 14, 14 body parts, respectively, for |*log*_2_(*FC*)|≥0.5, 1, 1.5, 2, 2.5). All comparisons between the median number of body parts influenced by *conserved* genes (median of 30 body parts influenced) and by *cis* or *cis+trans* genes at the various |*log*_2_(*FC*)| cutoffs are statistically significant (adjusted *p* < 0.04 by two-tailed Mann-Whitney U test after FDR correction). This trend does not hold for gene expression differences due to trans-regulatory changes (adjusted *p* > 0.19 for all comparisons between *conserved* genes and *trans* genes at the various |*log*_2_(*FC*)| cutoffs).

**Fig. S4.**
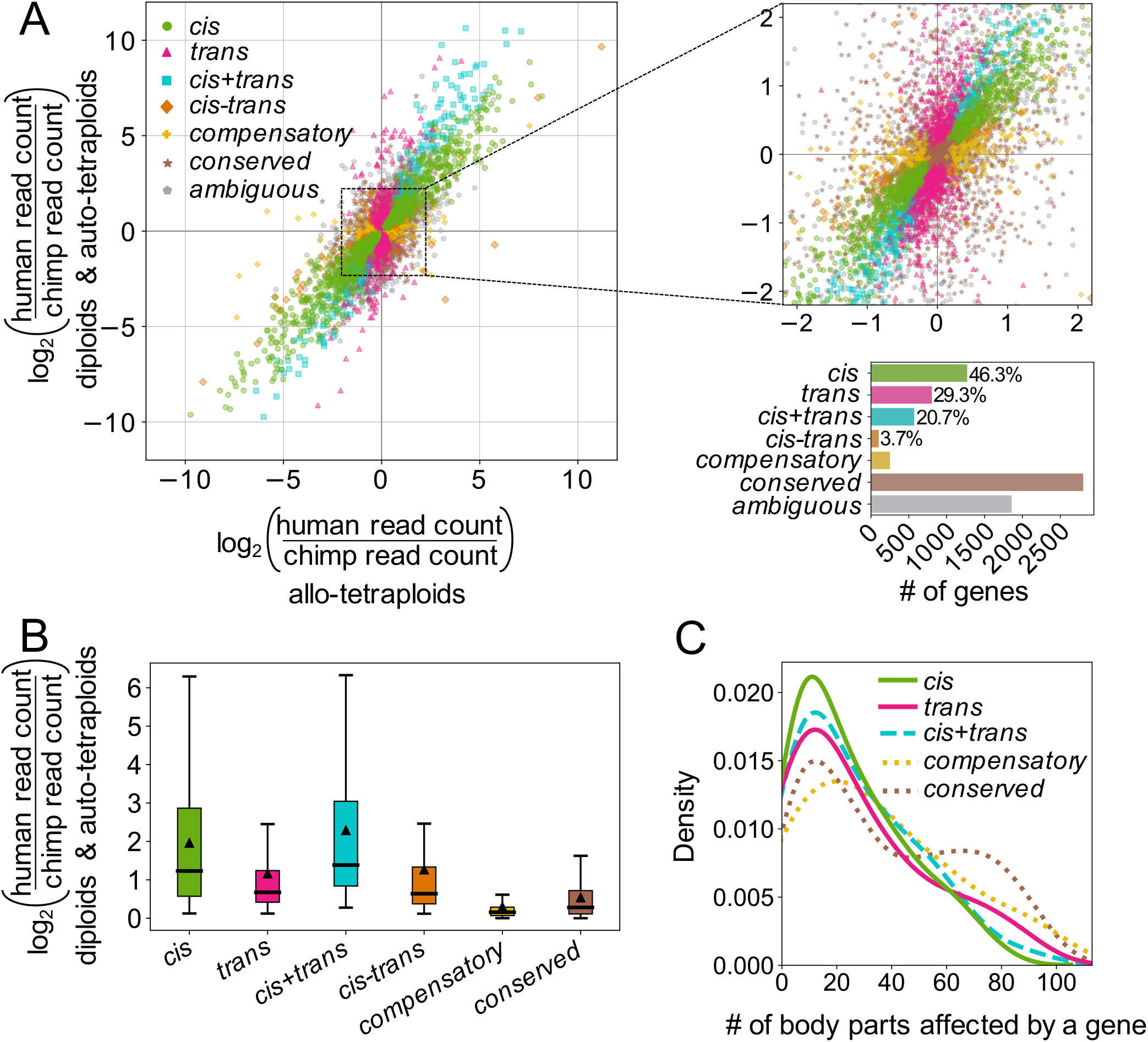
*Cis* and *trans* analysis results are robust to aneuploidies. Removing chromosomes with aneuploidies or abnormalities in any of the cell lines used for RNAseq does not meaningfully change the observed *cis* and *trans* trends demonstrated in Fig. 2C-D. In addition to genes on human chromosome 18 and their orthologous genes in chimpanzee (deletion of one copy of human chr18q is shared by a subset of cell lines and was removed for the *cis* and *trans* analysis shown in Fig. 2C-E), genes on human chromosomes 7, 12, 20, and Y (and orthologous genes in chimpanzee) and chimpanzee chromosomes 1, 2A, 2B, 11, 13, 14, 17, 19, 20, and Y (and orthologous genes in human) were removed prior to analysis. **(A)** See Fig. 2C legend. **(B)** See Fig. 2D legend. All pairwise comparisons are statistically significant by two-tailed Mann-Whitney U test after FDR correction with adjusted *p* < 10^−5^ except *trans* compared to *cis-trans* (*p* = 0.28). **(C)** See Fig. 2E legend. Genes classified as *cis, trans*, or *cis+trans* tend to influence fewer body parts than *conserved* genes (median 19, 20, 19.5 body parts, respectively, compared to median 29 body parts for *conserved* genes, adjusted *p* = 0.0029, 0.024, 0.024 by two-tailed Mann-Whitney U test after FDR correction). Note that the comparison between the *trans* and *conserved* categories is not statistically significant in Fig. 2E.

**Fig. S5.**
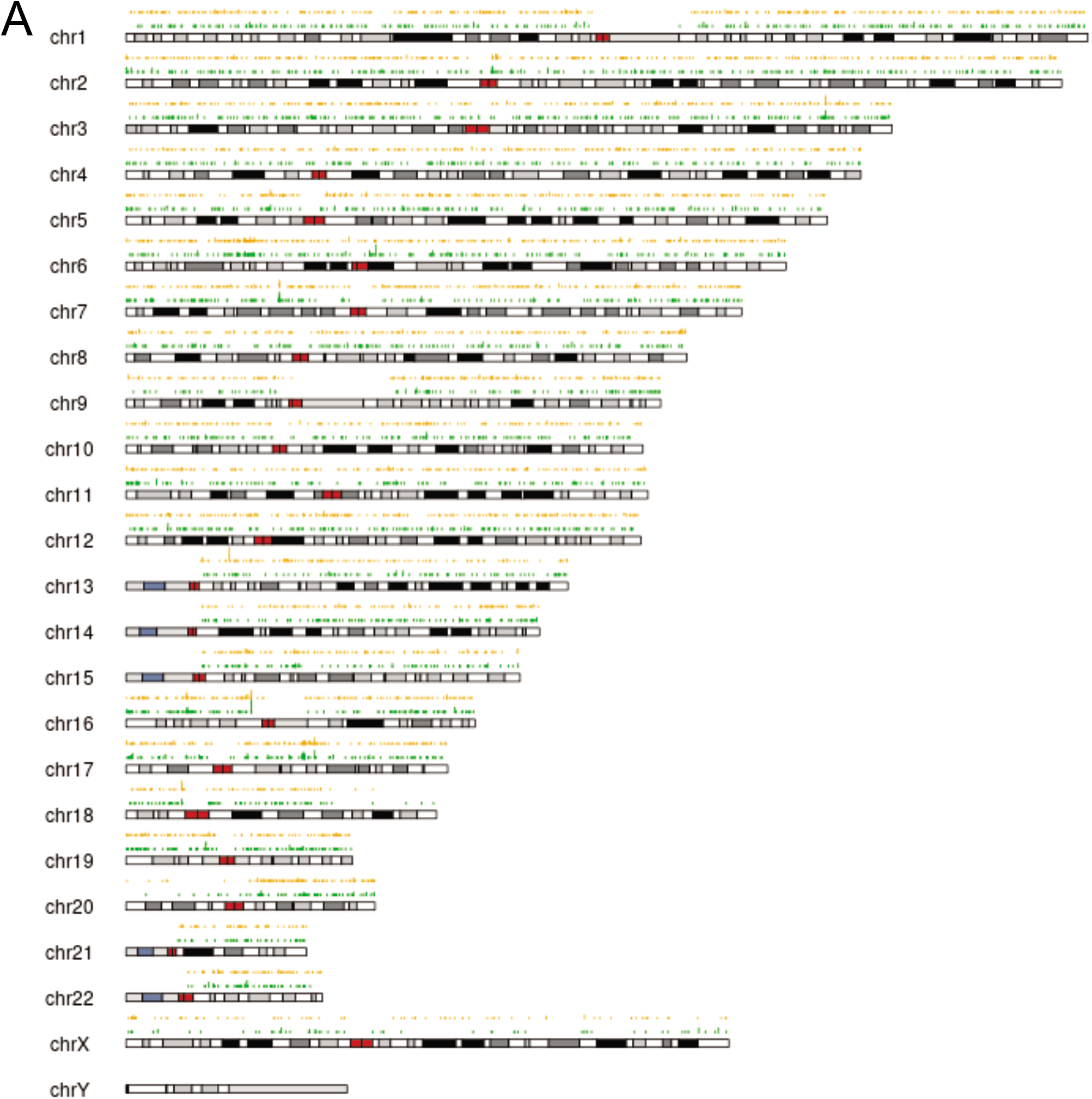
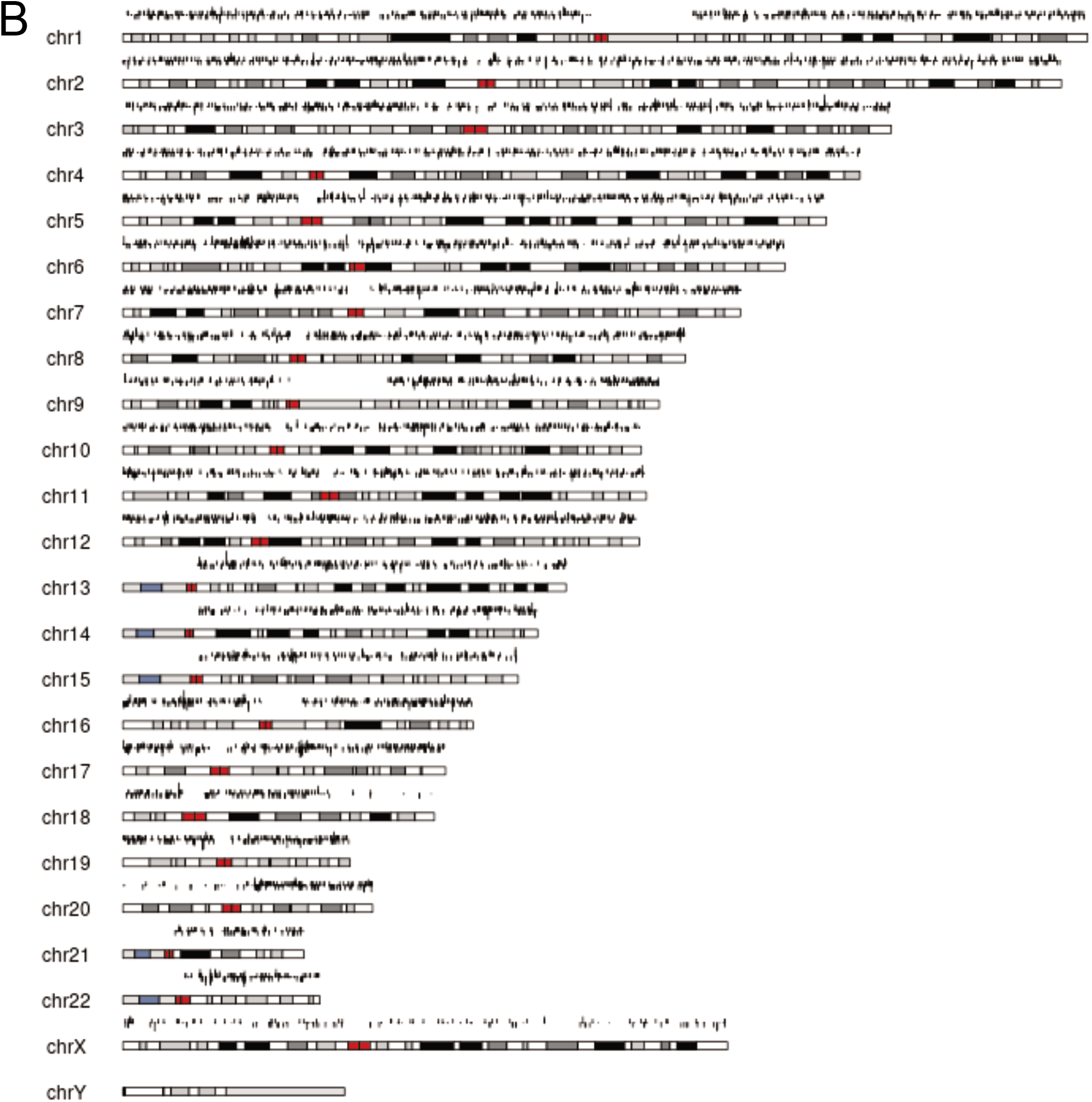
Distribution of genome-wide inter-specific recombination events identified by haplotagging. Representative chromosome plots showing locations of inter-specific molecules detected by haplotagging after genome-wide sequencing and filtering. **(A)** Comparison of inter-specific recombination events after different treatments. Green: cells treated with gRNA-chr20 (g20). Orange: cells treated with gRNA-chr20 and ML216 (g20+ML216). **(B)** *Log*_2_ ratio between the number of inter-specific recombination events along 10kb windows in g20+ML216 and g20 treated cells.

**Fig. S6.**
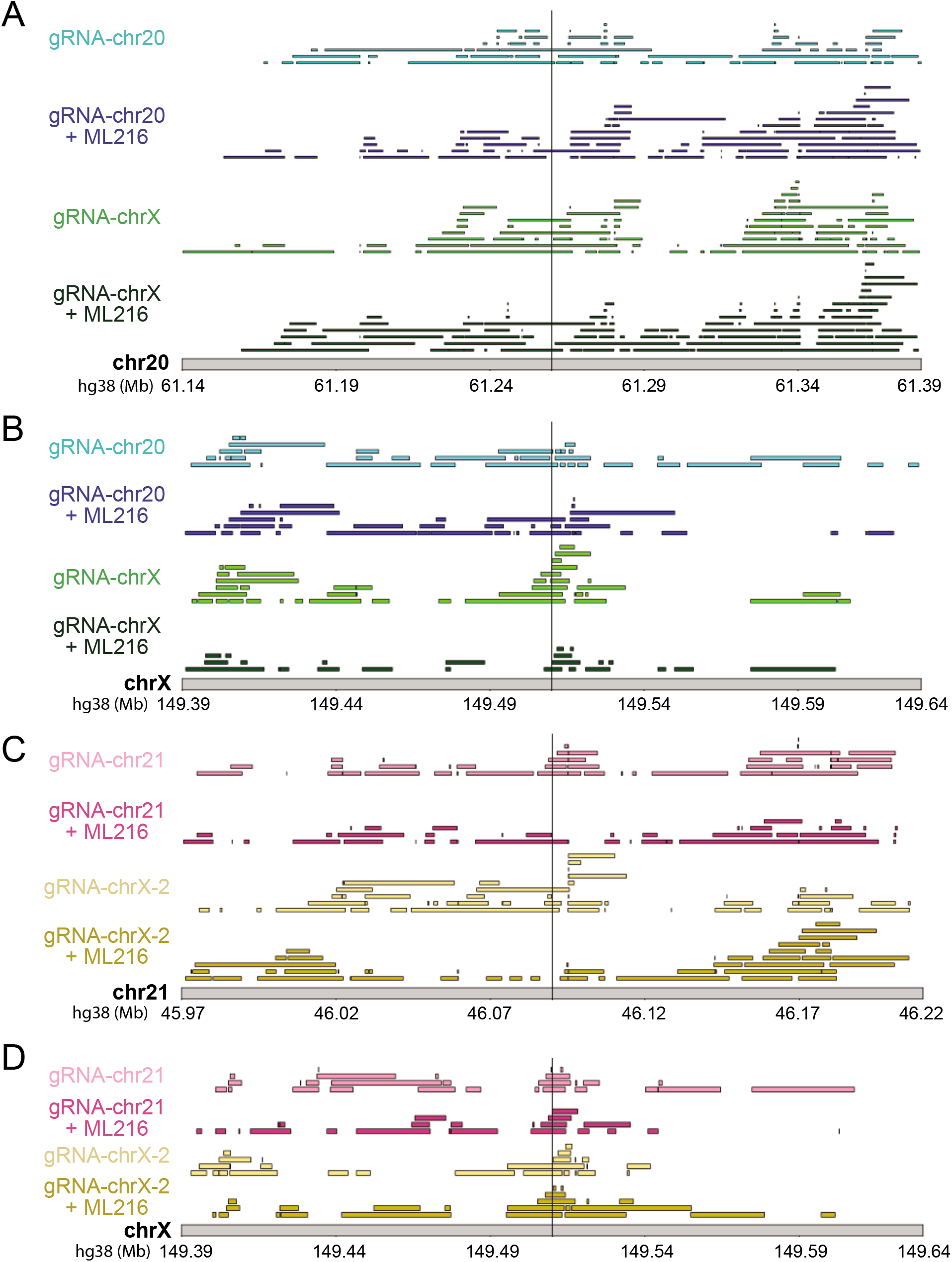
CRISPR targeting does not elevate inter-specific recombination rates at target loci. Plot of inter-specific recombination events in the 250 kb window surrounding CRISPR target loci on chr20 **(A)**, chrX **(B)**, chr21 **(C)**, or a second guide location on chrX **(D)**. Each horizontal rectangle represents the boundaries of an inter-specific recombination event detected by haplotagging. Vertical lines indicate the gRNA target site. Events are filtered for molecules that contain only 1 inter-specific event and have at least 2 supporting variants per species but are otherwise pre-filtering. The lack of enrichment at the target sites does not change with different filters (SI Methods).

**Fig. S7.**
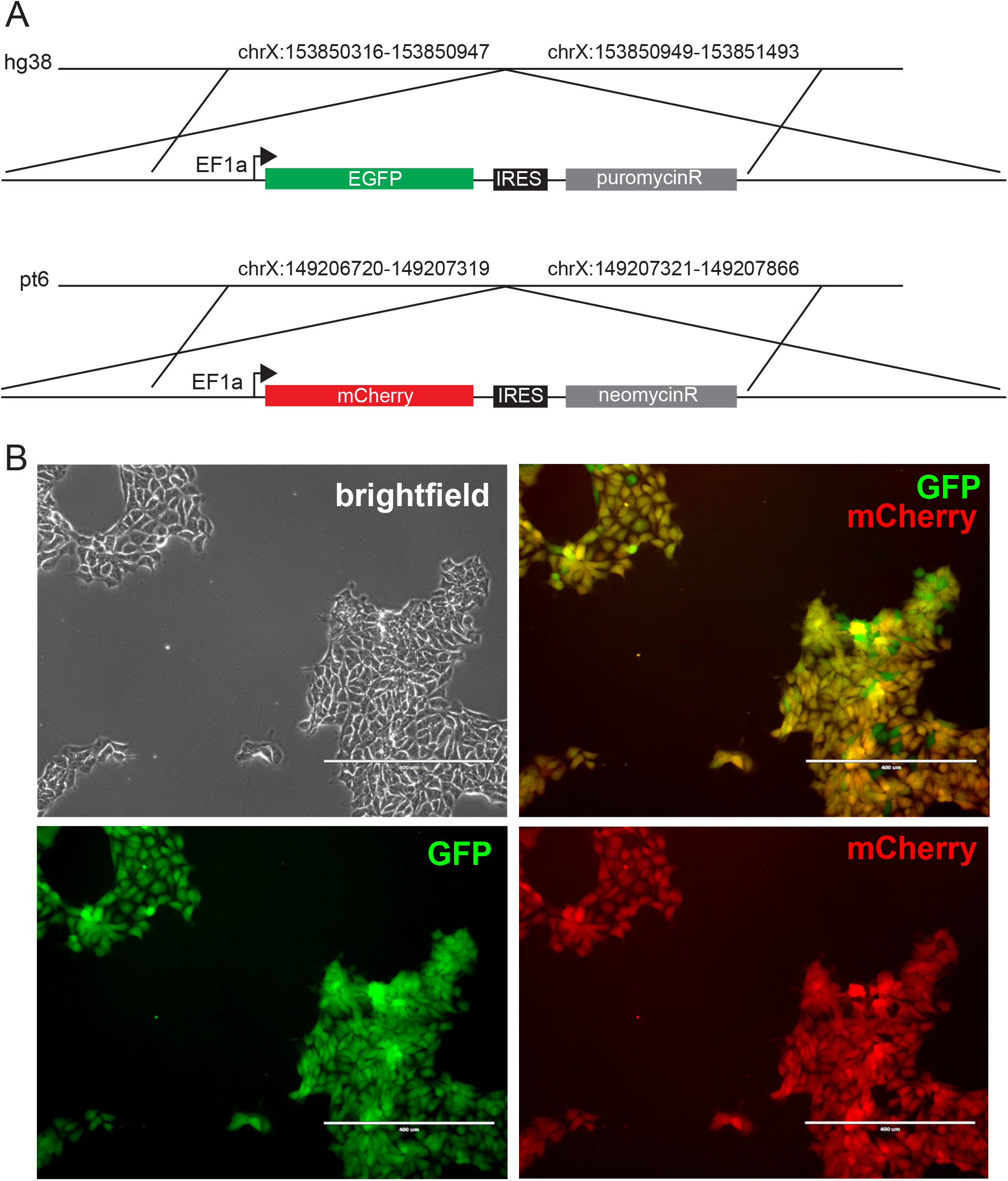
Generation of fluorescently-marked allo-tetraploid lines. Construct diagram and microscopy images for fluorescently-marked line. **(A)** Constructs containing EGFP or mCherry were inserted onto the human or chimpanzee chrX, respectively, using CRISPR-guided homologous recombination (Materials and Methods). Coordinates show locations of human and chimpanzee homology arms used in the constructs. **(B)** Allo-tetraploid H1C1a-X1 shown in brightfield, GFP, and mCherry. Cells marked with both GFP and mCherry appear yellow.

**Fig. S8.**
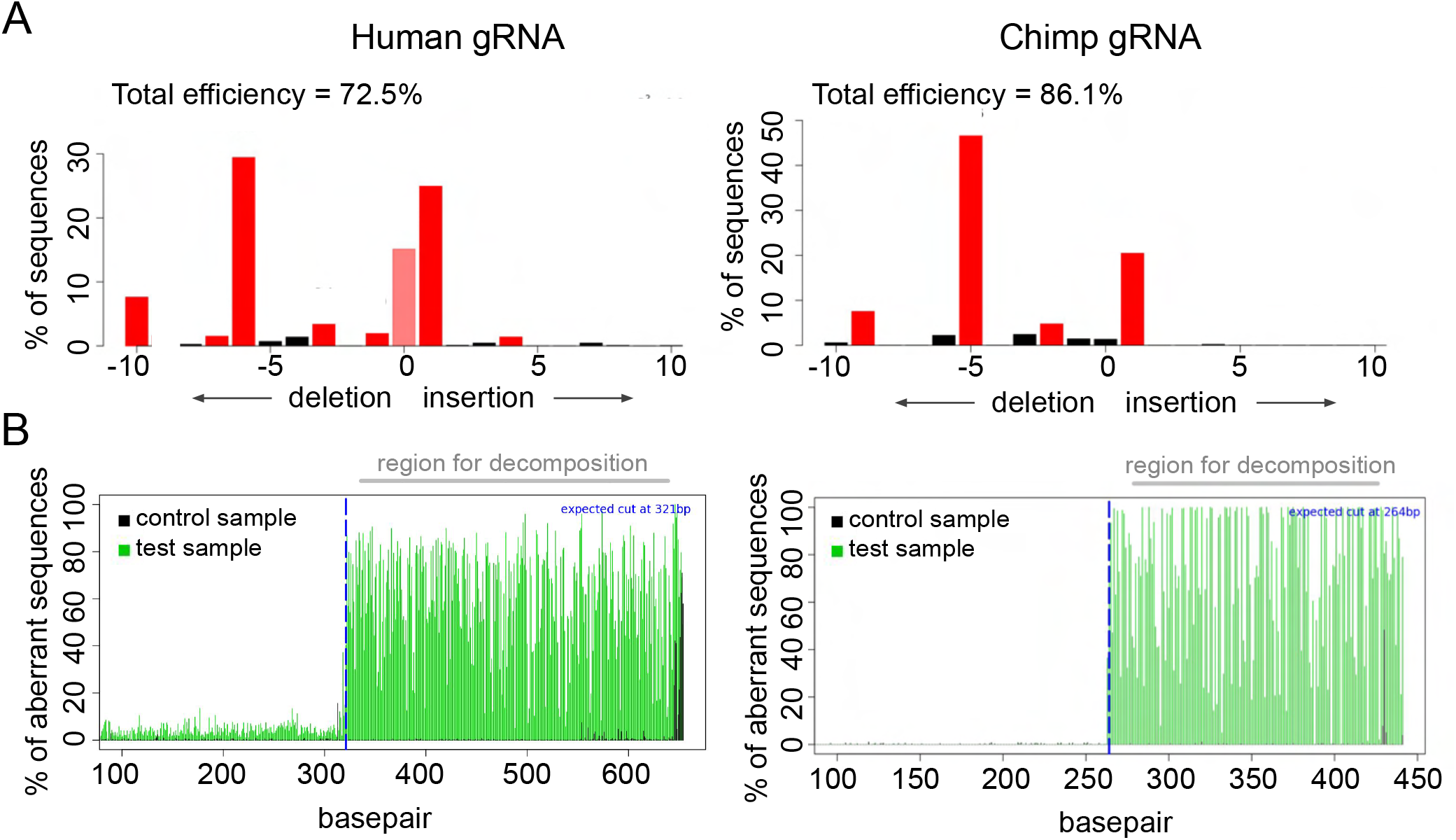
CRISPR/Cas9 gRNA editing efficiency and indel spectra for human and chimpanzee chrX guides. **(A)** For human- and chimpanzee-specific gRNAs, the spectrum and frequency of small insertions and deletions, gRNA efficiency, and **(B)** aberrant sequence signal plots are shown. Plots generated with Sanger sequence data in TIDE (Tracking of Indels by DEcomposition) (33).

**Fig. S9.**
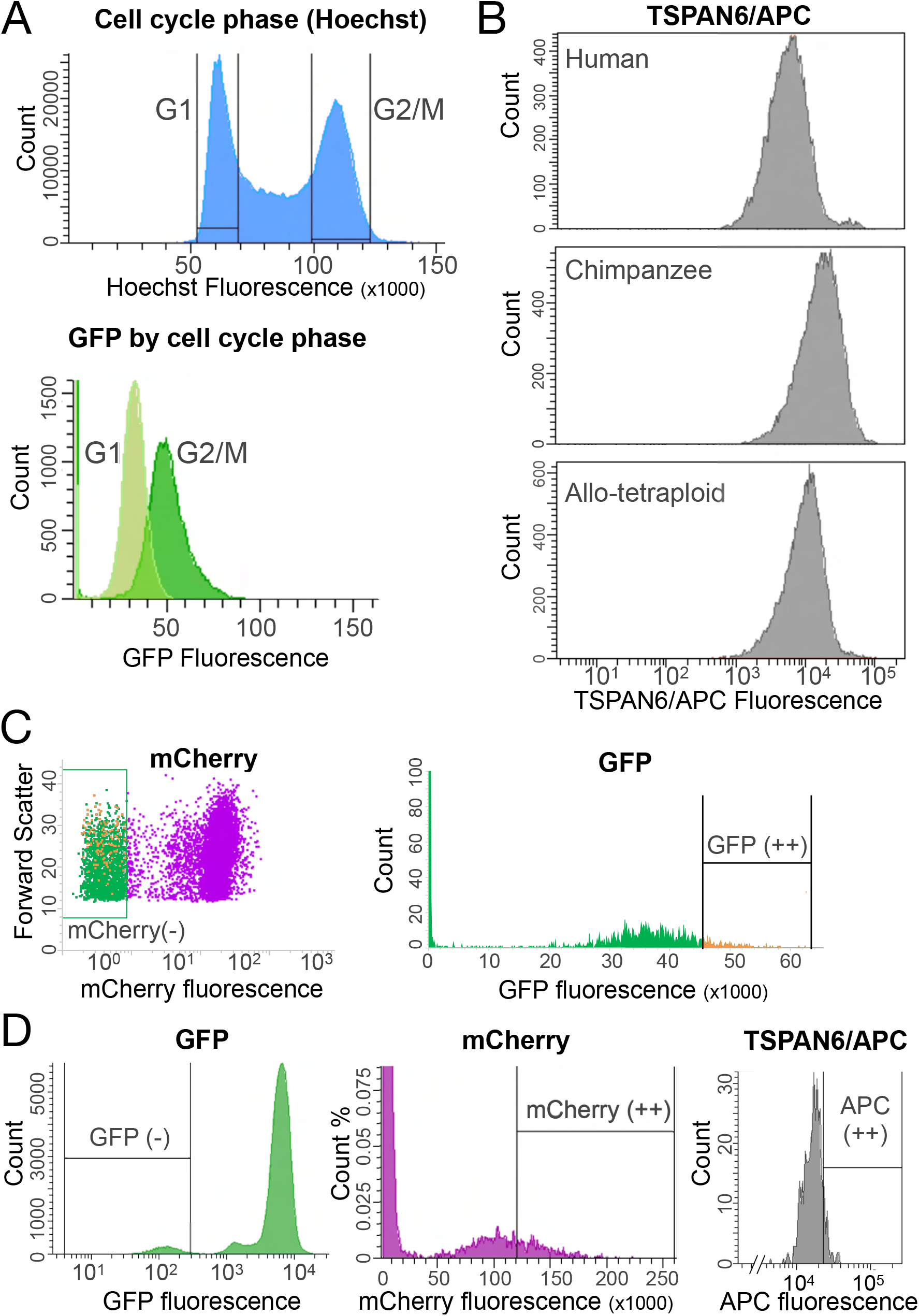
Fluorescence activated cell sorting (FACS) plots for chrX targeting. **(A)** Cell cycle phase determined by Hoechst peaks shows that G2/M cells exhibit higher GFP fluorescence than G1 cells. **(B)** Staining for TSPAN6 cell-surface protein with APC secondary antibody shows that chimpanzee TSPAN6-APC fluorescence intensity is higher than human TSPAN6-APC fluorescence intensity, with allo-tetraploid cells intermediate between human and chimpanzee values. **(C)** After G1 gating, cells treated with chimpanzee-specific gRNA are sorted for negative mCherry and high GFP fluorescence. **(D)** Cells treated with human-specific gRNA are sorted for negative GFP, high mCherry, and high TSPAN6-APC.

**Fig. S10.**
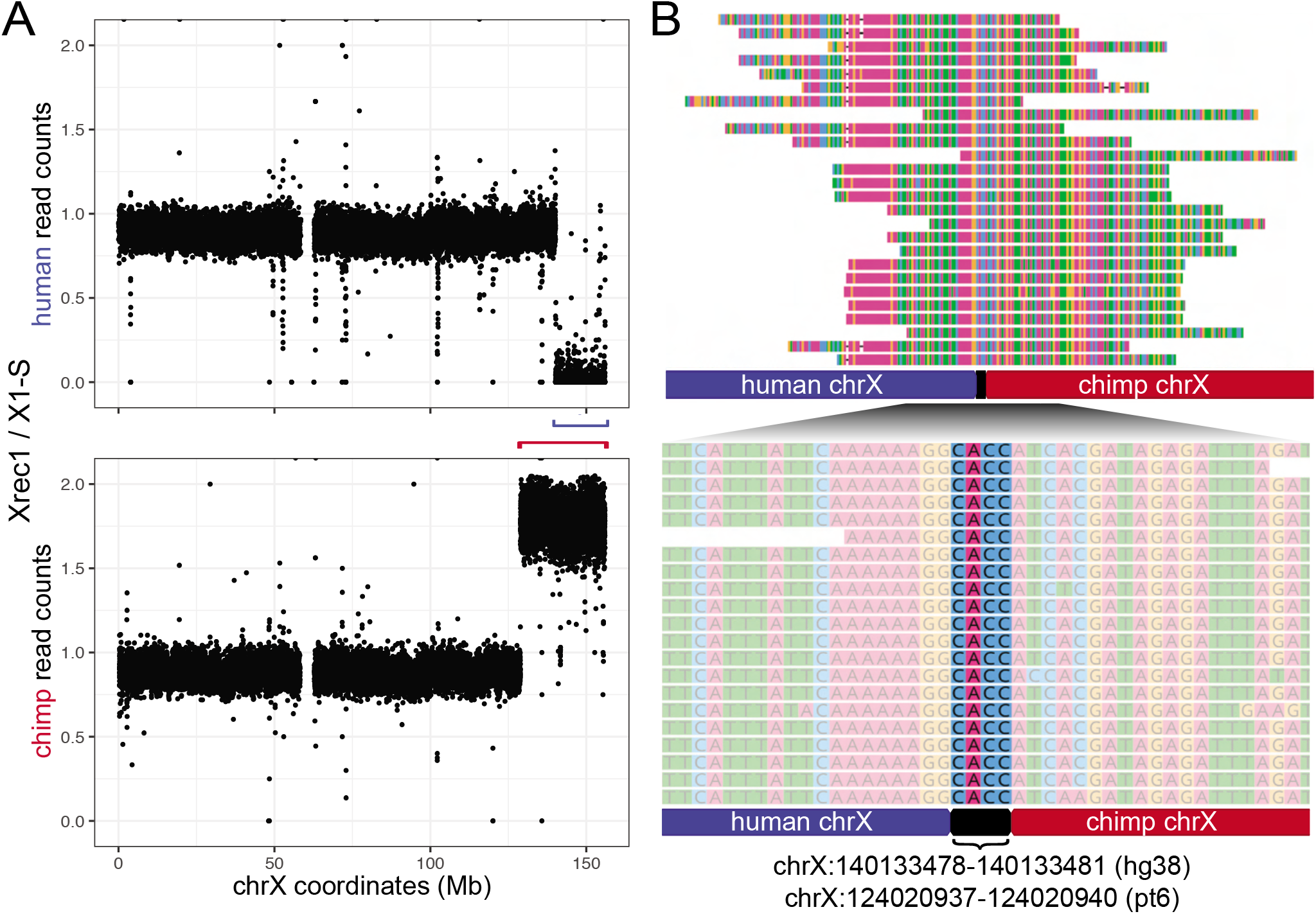
DNA sequencing identifies the site of recombination between human and chimpanzee X chromosomes. Whole-genome DNA sequencing data from the recombinant allo-tetraploid line H1C1a-X1-Xrec1 (Xrec1). **(A)** Read counts that align to either the human or chimpanzee allele along chrX were normalized to read counts for H1C1a-X1-S (X1-S), a control sample also sequenced in parallel (see Materials and Methods). This ratio was plotted along the X chromosome in hg38 coordinates. Blue bracket: region with no human read counts in Xrec1. Red bracket: larger region with twice as many chimpanzee read counts in Xrec1. **(B)** Reads that span the inter-specific recombination site in Xrec1 align to the appropriate locations in human chrX and chimpanzee chrX. The recombination site is a 4bp microhomology (highlighted region in close-up) that is found in both human chrX and chimpanzee chrX at the indicated coordinates.

**Fig. S11.**
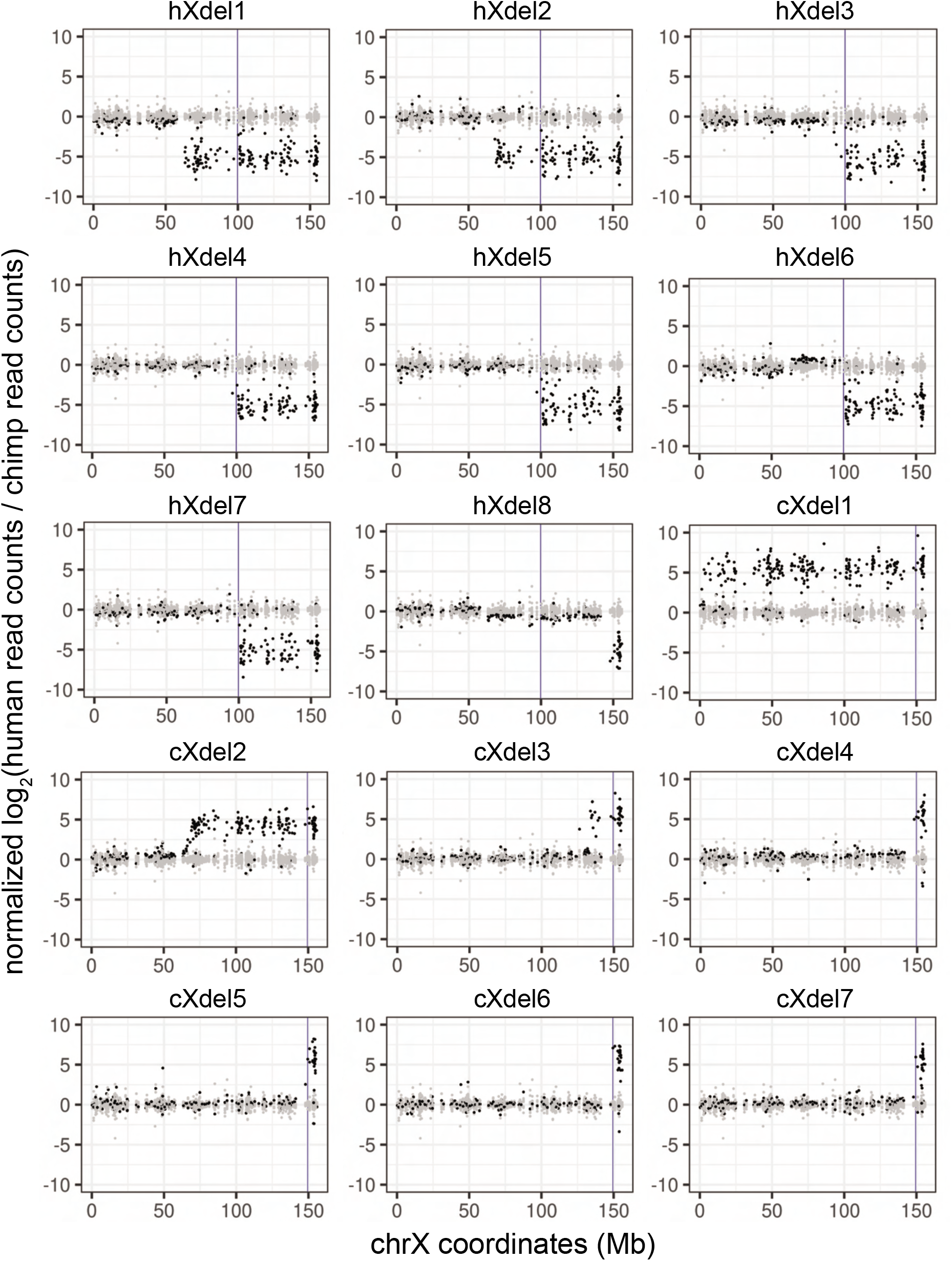
RNAseq of chrX lines localizes terminal deletion breakpoints. The relative allelic expression of genes along chromosome X is plotted for each of the chrX deletion lines (Table S1). The y-axis is the ratio of reads that map to the human or chimpanzee allele in the deletion line (black) normalized to the ratio of reads that map to the human or chimpanzee allele in the control (non-deletion) lines (gray) (see SI Methods). Each dot represents a gene on chromosome X plotted along the x-axis at its hg38 coordinate. The vertical line is the species-specific gRNA target site used to generate each deletion line.

**Fig. S12.**
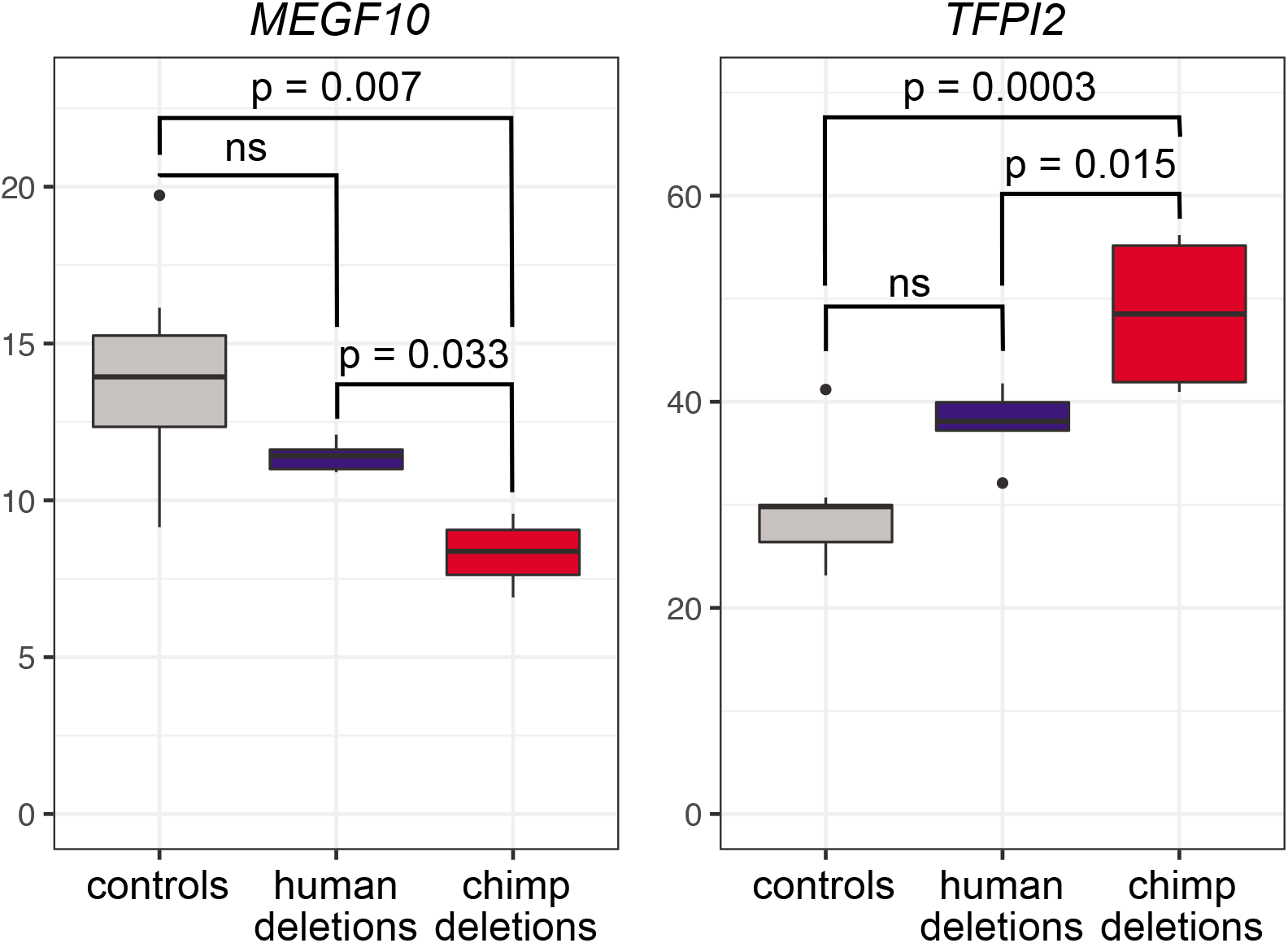
Differential expression of autosomal genes *MEGF10* and *TFPI2* in cell lines with distal chimpanzee chrX deletions compared to controls or to cells with distal human chrX deletions. Expression of *MEGF10* and *TFPI2* is significantly different in the four lines (cXdel4-cXdel7) with chimpanzee X chromosome deletions distal to breakpoints around 148Mb (in terms of hg38 coordinates) when compared to the nine control lines without deletions or to the five lines (hXdel3-hXdel7) with human X chromosome deletions distal to breakpoints around 95Mb (SI Methods). In contrast, there is not a significant difference in expression between the control lines and hXdel3-hXdel7.

**Fig. S13.**
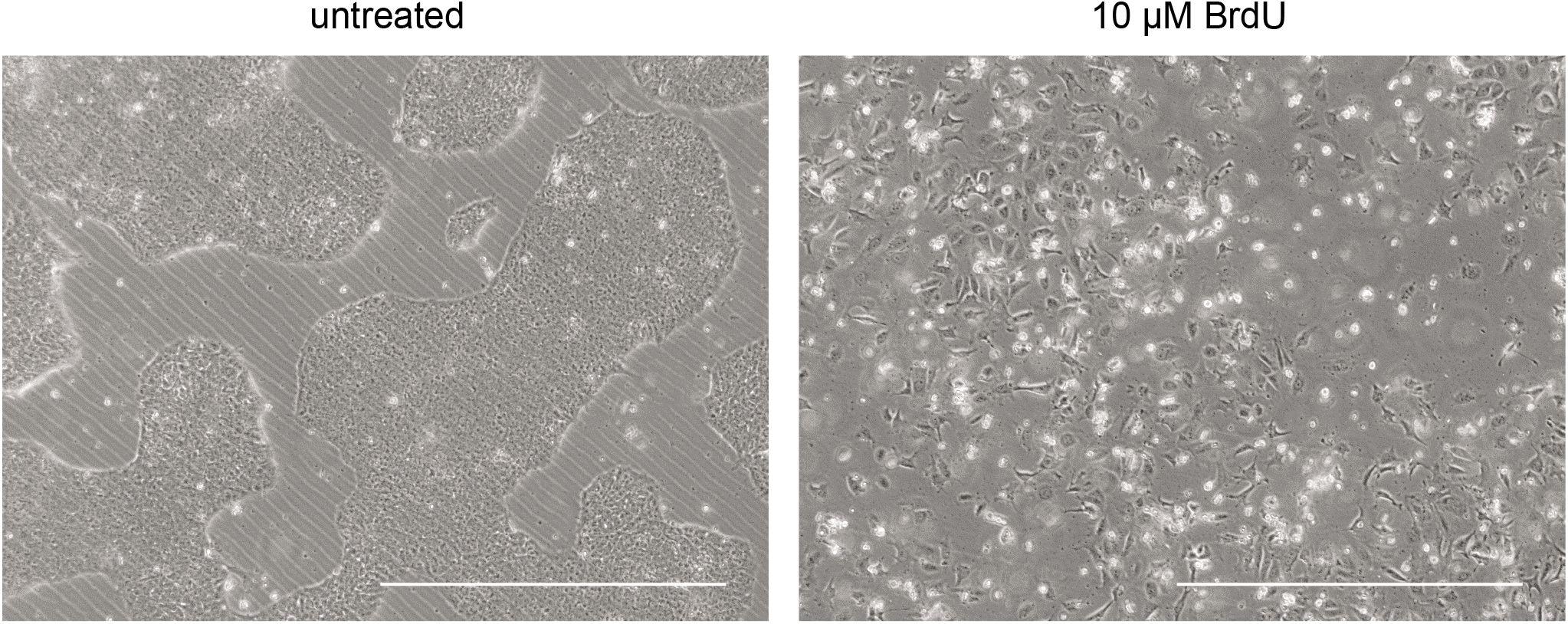
BrdU induces differentiation of iPSCs. After passaging, iPSCs treated with 10*μ*M of BrdU (right panel) are flatter and more spread out compared to untreated iPSCs (left panel). BrdU-treated cells also do not form the colonies typical of iPSCs and fail to divide, suggesting that they have terminally differentiated. Scale bars are 1mm.

**SI Dataset S1 (Table S1)**

iPSC lines used and generated in the current study.

**SI Dataset S2 (Table S2)**

Primers and gRNAs.

**SI Dataset S3 (Table S3)**

Trilineage differentiation results.

**SI Dataset S4 (Table S4)**

Differential gene expression analysis of diploid and auto-tetraploid iPSC lines.

**SI Dataset S5 (Table S5)**

Differential gene expression, allele-specific gene expression, and regulatory type (*cis/trans*) analysis between humans and chimpanzees in diploid, auto-tetraploid, and allo-tetraploid iPSC lines.

**SI Dataset S6 (Table S6)**

Gene ontology enrichments for regulatory type (*cis/trans*) categories.

**SI Dataset S7 (Table S7)**

qPCR and PCR results on chrX for sorted colonies treated with CRISPR+ML216.

**SI Dataset S8 (Table S8)**

Differential gene expression analysis of chrX deletion iPSC lines.

## Notes

The authors declare no competing interests.

### Competing Interest Statement

The authors have declared no competing interest.

